# Social and environmental transmission spread different sets of gut microbes in wild mice

**DOI:** 10.1101/2023.07.20.549849

**Authors:** Aura Raulo, Paul Bürkner, Jarrah Dale, Holly English, Genevieve Finerty, Curt Lamberth, Josh A Firth, Tim Coulson, Sarah CL Knowles

## Abstract

Gut microbes shape many aspects of organismal biology, yet how these key bacteria transmit among hosts in natural populations remains poorly understood. Recent work in mammals has emphasized either transmission through social contacts or indirect transmission through environmental contact, but the relative importance of different routes has not been directly assessed. Here, we used a novel RFID-based tracking system to collect long-term high resolution data on social relationships, space use and microhabitat in a wild population of mice (*Apodemus sylvaticus*), while regularly characterising their gut microbiota. Through probabilistic modelling of the resulting data, we identify positive and statistically distinct signals of social and environmental transmission, captured by social networks and overlap in home ranges respectively. Strikingly, microbes with distinct biological attributes drove these different transmission signals. While aerotolerant spore-forming bacteria drove the effect of shared space use, a mix of taxa but especially anaerobic bacteria underpinned the social network’s effect on gut microbiota similarity. These findings provide the first evidence for parallel social and environmental transmission of gut microbes that involve biologically distinct subsets of the mammalian gut microbiota.

**List of contributions:**

- *Aura Raulo* designed the study, helped develop the new RFID tracking technology, collected the data from Wytham, completed all laboratory analyses on gut microbiota profiling prior to sequencing, developed analytical methods, analysed the data and wrote the manuscript
- *Paul Bürkner* helped design the Bayesian probabilistic modeling framework and provided feedback on the manuscript
- *Jarrah Dale* helped collect field data using RFID loggers
- *Holly English* helped collect field data using RFID loggers and provided feedback on home range analyses
- *Genevieve Finerty* helped with home range analysis and the analysis of microhabitat variation and provided feedback on the manuscript
- *Curt Lamberth* led development of RFID tracking devices and helped collect field data from Wytham
- *Josh Firth* supervised the research project, developed social network analysis methods and provided feedback on the analyses and the manuscript
- *Tim Coulson* supervised the research project and provided feedback on the analyses and the manuscript
- *Sarah Knowles* supervised the research project, helped develop the tracking technology and design the study, collected data from Wytham, planned and supervised laboratory methods, developed analytical methods and provided feedback on analyses and the manuscript.

## Introduction

Host-associated microbiotas, especially the diverse communities inhabiting the vertebrate gut, are increasingly recognised as key influencers of their host’s biology, affecting the development (1–3), physiology (4,5), behaviour (6–8) and ultimately ecology and evolution of their host (9–12). Many biological effects of the microbiota depend on community composition, which can show vast multidimensional variation among individuals, populations and species, as well as strong temporal dynamics within individuals.

Although gut microbiota variation is thought to have important effects on animal fitness, our understanding of how different processes come together to shape microbiota in natural populations remains limited. As with any ecological community, fundamental ecological processes will govern community assembly of the microbiota (13–15). These include processes operating inside hosts, such as microbe-microbe and host-microbe interactions (10), but importantly also processes operating outside the host, which affect how microbes come to colonise hosts in the first place. Host-associated microbes live an inherently patchy landscape, with hosts forming habitat islands in a sea of less suitable habitat through which they must disperse. As such, microbiotas are well conceptualised as metacommunities (16) whereby microbial transmission from other hosts and the environment constitute potentially powerful forces shaping composition of individual microbiotas (17).

Gut microbes can colonise hosts through various routes. In mammals, transmission starts at birth with colonisation by microbes in the birth canal and from the mother’s gut microbiota (18), and continues throughout life as microbes spread through contacts with conspecifics as well as the wider ecosystem. Recent research has specifically emphasized the importance of animal social behavior in the spread of gut microbes (19). Host-to-host transmission can occur either via direct contact during social behaviours, or indirectly through host microbial shedding to and acquisition from a shared environment. The host social network has been framed as an important microbial transmission landscape, “a social archipelago of host islands” that shapes microbial community structure (19). In humans, sharing a living space predicts sharing of gut microbes (20–24), typically much more so than genetic relatedness (21,24). The intimacy of social interaction also appears to be important, with friends and spouses sharing more gut microbes than strangers, with the effect strongest among spouses self-reporting a physically close relationship (25). Social group membership in other group-living mammals also predicts gut microbiota composition (26–34) and within social groups, stronger pairwise social relationships can predict a higher degree of microbiota similarity (26–28). Such effects have also been recently documented in less social species that do not form social groups. In wild wood mice, we recently showed that social networks predicted the sharing of gut microbes more strongly than genetic relatedness, seasonality and spatial proximity (35).

A separate body of research has emphasised that microbiotas can also be shaped by contact with the broader natural environment, such as soil and food. For example, the gut and skin microbiota of human children was shown to be markedly influenced by variable physical contact with local biodiversity and natural soils (36–39), and experimental soil exposure can change the gut microbiota of laboratory mice (40,41). Gut and skin microbes have also been shown to spread between humans through their shared environment. For example, sharing a room was associated with gradual homogenisation of microbiota between students paralleled with microbiota of the room surfaces becoming gradually more similar to that of the people living in it (42). A recent study also found that human gut microbes can persist on built environment surfaces long enough to be transmitted between people: majority of viable human-derived microbiota detected on bathroom floors were aerobic taxa, though some methanogenic anaerobes could also stay alive on building surfaces for up to 6 hours (43). Studies such as these challenge the idea of microbiomes as strict metacommunities (akin to oceanic island systems), as some gut-dwelling microbes clearly persist and perhaps even flourish outside hosts (42–46).

Despite evidence that both social interaction and environmental transmission can shape vertebrate gut microbiotas, the influence of these different routes have rarely been studied together and directly compared. For example, although environmental exposure (the degree of contact with natural soils and local environments) has been implicated in driving healthy microbiota development and immune function in human children (36–39), the role of socially transmitted bacteria in the same processes has not been examined. The opposite bias prevails in studies of social transmission, with most non-human studies ignoring the potential impact of environmental transmission (but see (47)). Consequently, we know little about the relative importance of social and environmental transmission in shaping the gut microbiota.

Importantly, bacteria vary in many attributes that may affect their propensity to transmit via different routes. Traits that influence their ability to persist and grow outside the host, such as aerotolerance and spore formation, may be particularly important in this regard (48). For example, aerotolerant microbes may persist and even grow outside terrestrial host organisms (49), spore-formers may be able to persist long enough outside hosts to transmit indirectly via environmental contact (50), while anaerobic, non-spore forming bacteria may instead rely on intimate physical contact to pass from host to host (51). Consistent with such ideas, among Firmicutes spore-forming taxa are more prevalent among humans than non-spore-forming taxa, implying they may be more readily transmitted through shared environments (52). Aerotolerant and spore-forming bacteria in the human gut were also found to have broader geographic range, suggesting they can spread across larger distances than those less tolerant of oxygen-rich environments (21). However, to date no empirical work has formerly tested whether gut bacterial phenotypes predict which transmission routes are most responsible for spreading them.

To bridge this gap, here we use a tractable wild mammal system to dissect how both social and environmental transmission shape gut microbiota composition, and ask which types of microbial taxa are shared via each route. Delineating separate signals of social and environmental transmission can be challenging in some species including humans (53), particularly those that form tight social groups where social interactions, spatial location and other factors that shape the microbiota (such as diet) are highly correlated. We therefore opted to use a semi-social model species, the wood mouse (*Apodemus sylvaticus*) for this purpose. These nocturnal woodland rodents inhabit small, stable home ranges and have non-modular social networks in which social relationships are only partially related to the sharing of space (35), making them well-suited to disentangling social and environmental transmission effects on microbiota composition. In this species, social transmission of microbes could occur through physical contact behaviours (e.g., mating, huddling, grooming, licking, fighting, see Supplementary Figure S1), whereas environmental transmission could happen through contact with microbes present in natural surfaces, soil, food items or other’s faeces (though coprophagy behaviour has not been observed in this species). Within a single population over a 10-month period, we used a passive tracking system based on radio-frequency identification (RFID) technology to intensively monitor home ranges, social networks and microhabitat use, while in parallel repeatedly profiling individuals’ gut microbiota from faecal samples. With the resultant data, we then dissect how sharing of gut microbial taxa among individuals varies as a function of their social association, overlap in space use, and similarity in habitat, and how microbial traits (aerotolerance and spore-forming ability) predict the extent to which microbes drive distinct transmission signals.

## Results

### The wood mouse gut microbiota is highly individualised and temporally variable

We first explored the taxonomic composition of wood mouse gut microbiota, which we found to be dominated by bacteria belonging to the families Lachnospiraceae (37% amplicon sequence variants, hereafter ASVs), Muribaculaceae, (formerly known as S24-7; 20% ASVs), Oscillopiraceae (8% ASVs) and Ruminococcaceae (4% ASVs). The most common genera were *Lactobacillus*, Lachnospiraceae NK4A136, *Ligilactobacillus* and *Limosilactobacillus* (Supplementary Figure S2A). Using a set of repeat-sampled mice (mice with ≥2 samples, 255 samples from 82 individuals), we found microbiota composition to be highly individualised, with individual identity explaining 52% variation in community composition (marginal PERMANOVA on Jaccard Index, R^2^=0.52, F=2.38, p=0.001). Microbiota composition also varied temporally, with sampling month explaining 5% of compositional variation in the same analysis (R^2^= 0.05, F=1.67, p=0.001; Supplementary Figure S2B).

### Wood mice have weakly correlated social and spatial population structure

To identify different transmission pathways for microbiota, we used logger data to derive home ranges and microhabitat profiles for each individual mouse, and social networks for the population. Social networks were constructed using the “Adjusted Simple Ratio Index” (Adjusted SRI) as a measure of social association, which reflects how often two mice were observed in the same location within 12h of each other at times they were both alive, during a specified period of time (see Methods). Individual home ranges (space-utilisation distributions) were calculated from logger data and used to calculate pairwise home range overlap using the Bhattacaryya index. To measure whether mice were exposed to similar microhabitats, we used data from a ground-cover vegetation survey to calculate an index of vegetation community similarity (Bray-Curtis index) across each pairs home ranges. The mouse social network displayed a non-modular structure, with no clustering into social groups (Figure 1). Across the entire 10-month monitoring period, mice had a mean of 6.4 social connections (i.e. 6.4 other mice with which they occurred in the same location at least once within a 12 h window, range 0-24), though these varied considerably in association strength (mean non-zero social association index=0.10, sd=0.15, range=0.005-1). We also constructed separate social networks for spring (Feb-Jun) and Fall (Jul-Nov) to assess how social associations changed across the annual reproductive cycle and with the sex of pair members. In spring, average social association strength was comparable across different sex categories (female-female, female-male and male-male pairs) but tended to be stronger in spring compared to fall, especially in male-male pairs (Supplementary Figure S3A). Home range overlap also varied according to season and sex; in spring, home range overlap was greatest among males, intermediate among male-female pairs and lowest among females, while during Fall all sex-categories had similar levels of range overlap (Supplementary Figure S3A). Although habitat similarity varied widely across pairs of mice (mean Bray-Curtis habitat similarity=0.39, sd=0.16, range=0-0.98), it did not differ significantly between sex categories or seasons. Across the entire dataset, social association strength, spatial (home range) overlap and habitat similarity among mouse pairs were all partially correlated with each other, with the correlation between social networks and home range overlap much weaker than that between home range overlap and habitat similarity (Mantel tests: social association-spatial overlap, r=0.26, spatial overlap-habitat similarity r=0.50, Social association-habitat similarity r=0.12, all p<0.001, Supplementary Figure S3B).

**Figure 1.**
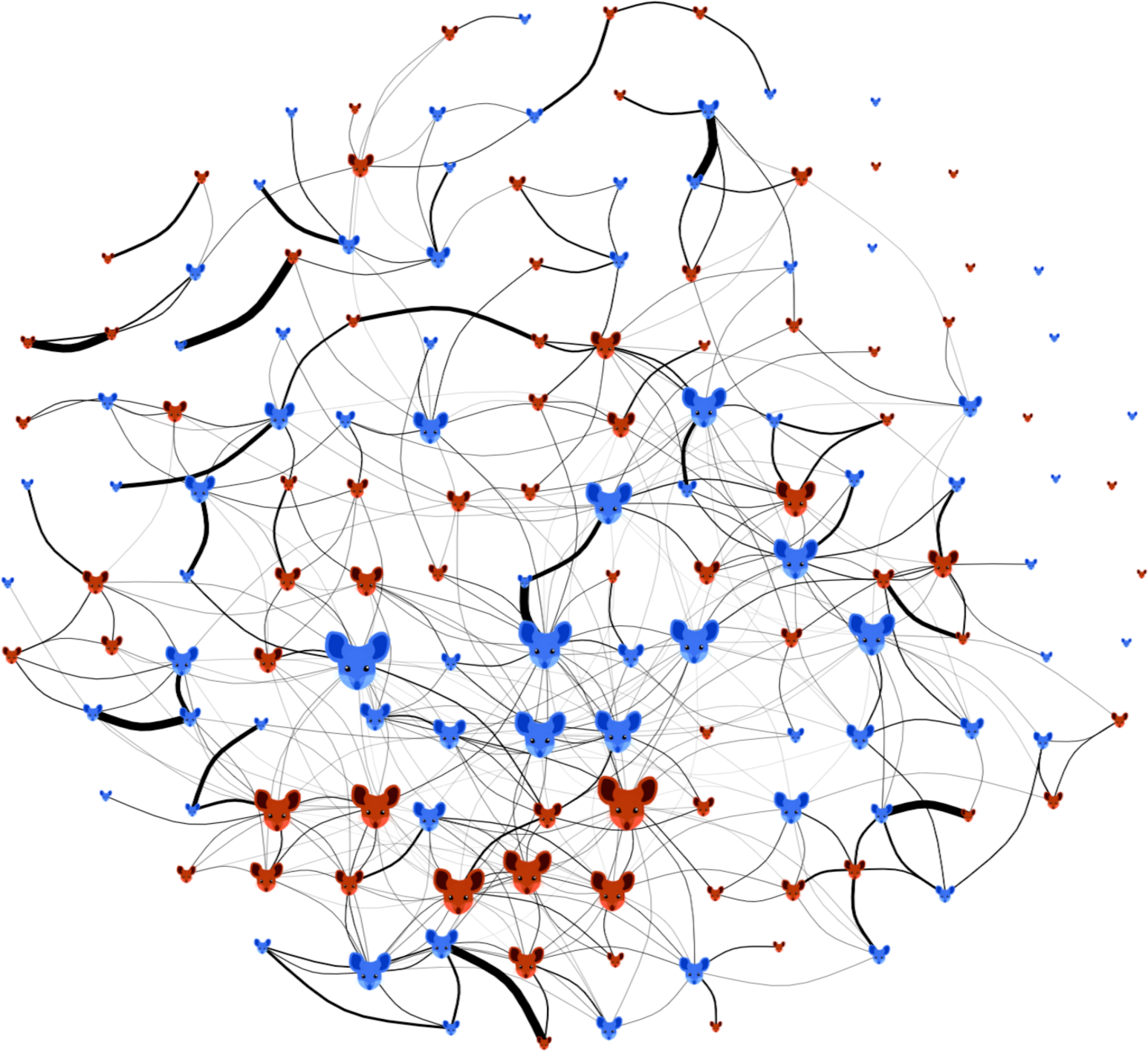
Social network of wood mice. Nodes are individual mice, either males (blue) or females (red). Edges are measures of social association (Adjusted SRI, see Methods). Node size reflects an individual’s degree i.e., the number of social connections (larger = more connections), and line thickness denotes social association strength (thicker = higher Adjusted SRI). Nodes are arranged in a standard spring layout forced into a circular fit.

### Social association, spatial overlap and habitat similarity predict gut microbiota similarity among mice

We constructed a dyadic Bayesian betaregression model to predict the level of microbiota similarity as a function of our measures of social association, spatial overlap and habitat similarity across all pairs of mice. Microbiota similarity was calculated as the proportion of shared 16S amplicon sequence variants (Jaccard index) among samples. Model results revealed that social association, spatial overlap and habitat similarity all positively predicted the proportion of microbial ASVs shared by pairs of mice, while controlling for other covariates (Figure 2). Social association had by far the strongest effect on gut microbe sharing – over eight times stronger than the effects of spatial overlap or habitat similarity (Figure 2, Supplementary Table S1A).

**Figure 2.**
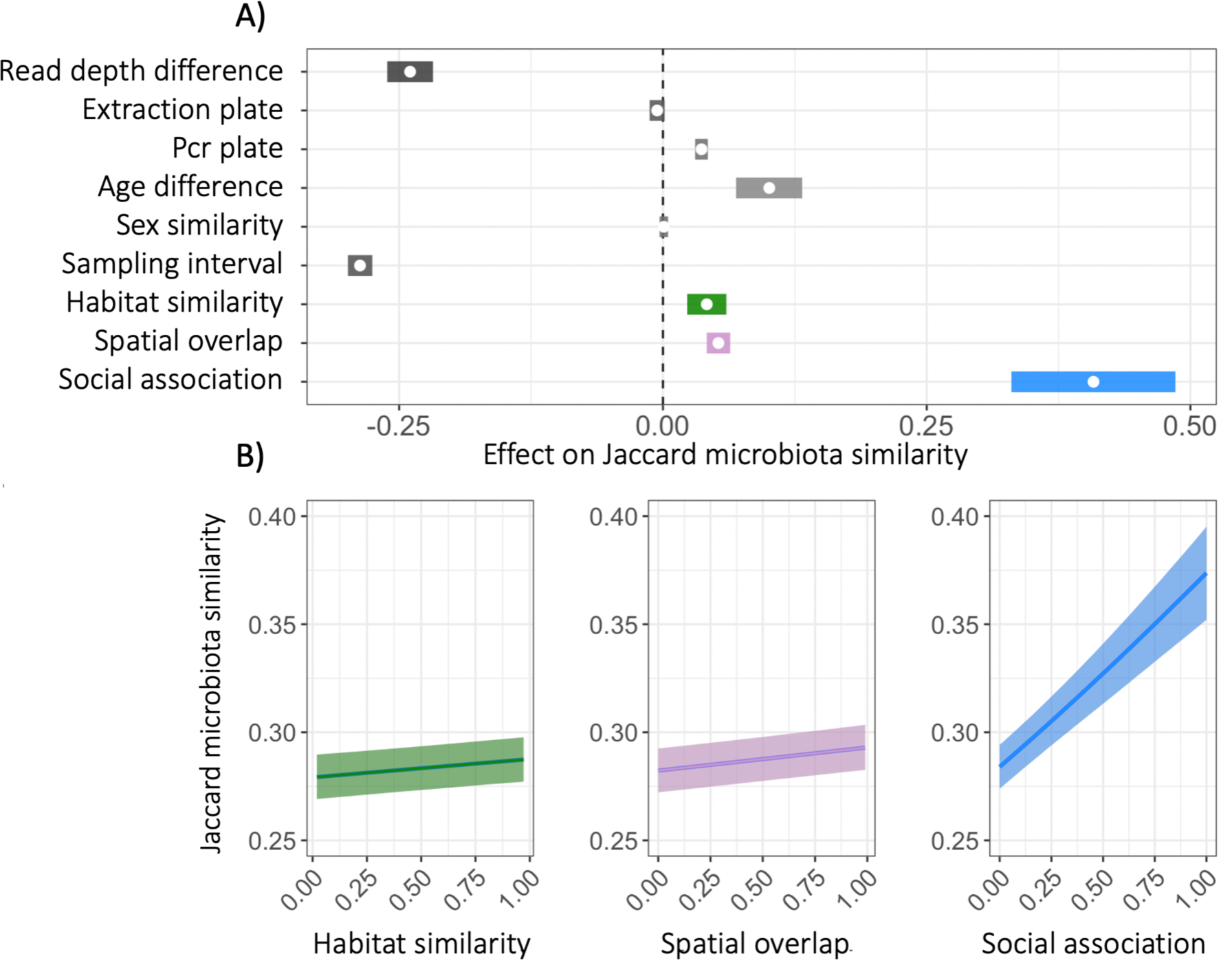
Effects of different predictors on microbiota similarity. A) Posterior means (points) and their 95% credible intervals (coloured lines) are plotted from Bayesian beta regression (brms) models (Supplementary Table S1A) with pairwise microbiota similarity among hosts (Jaccard Index) as the response. Where credible intervals do not overlap zero, a variable significantly predicts microbiota similarity while controlling for all other terms shown. B) Slopes of the three main predictors from the same model: habitat similarity (green), Spatial overlap (purple), social association (blue).

To further explore the nature of the social, spatial and habitat signals in the data, we ran the same model with two alternate response variables capturing different elements of microbiota similarity: Bray-Curtis similarity (1-Bray-Curtis distance) and the count of shared taxa between a sample pair, modelling using betaregression and a poisson model respectively. While effects were of similar magnitude in all models, social and spatial effects appeared notably more uncertain (displaying doubly as wide credible intervals) when using Bray-Curtis similarity compared to the Jaccard Index or count of shared taxa (Supplementary Figure S4, Supplementary Tables S1B-C).

We next explored whether the social, spatial and habitat effects on microbiota similarity varied according to the sex of individuals involved. For all three effects, we detected significant interactions with sex category (Supplementary Table S2), which showed that social association and home range overlap most strongly predicted gut microbe sharing among female-only pairs, while the habitat effect was strongest for male-male pairs (Figure 3). Because wood mouse social behaviour may be expected to vary across breeding and non-breeding seasons, we further explored whether the sex-dependent social association effect varied between spring and fall. This revealed that in the spring (Feb-Jun) the effect of social association on gut microbe sharing was only significant for female-only pairs, while in the fall (Jul-Nov), the social effect was driven by same-sex (both male-male and female-female) pairs (Supplementary Figure S5, Supplementary Table S3). Across all models, the effect of social association was strongest in female-only pairs during spring, where in real terms it meant that while female pairs that were never observed together shared on average 30% of their gut microbial taxa (ASVs), pairs with strong social associations (mice that were associated in over 50% of the instances they were observed) were predicted to share on average 60% of their combined gut microbial taxa.

**Figure 3.**
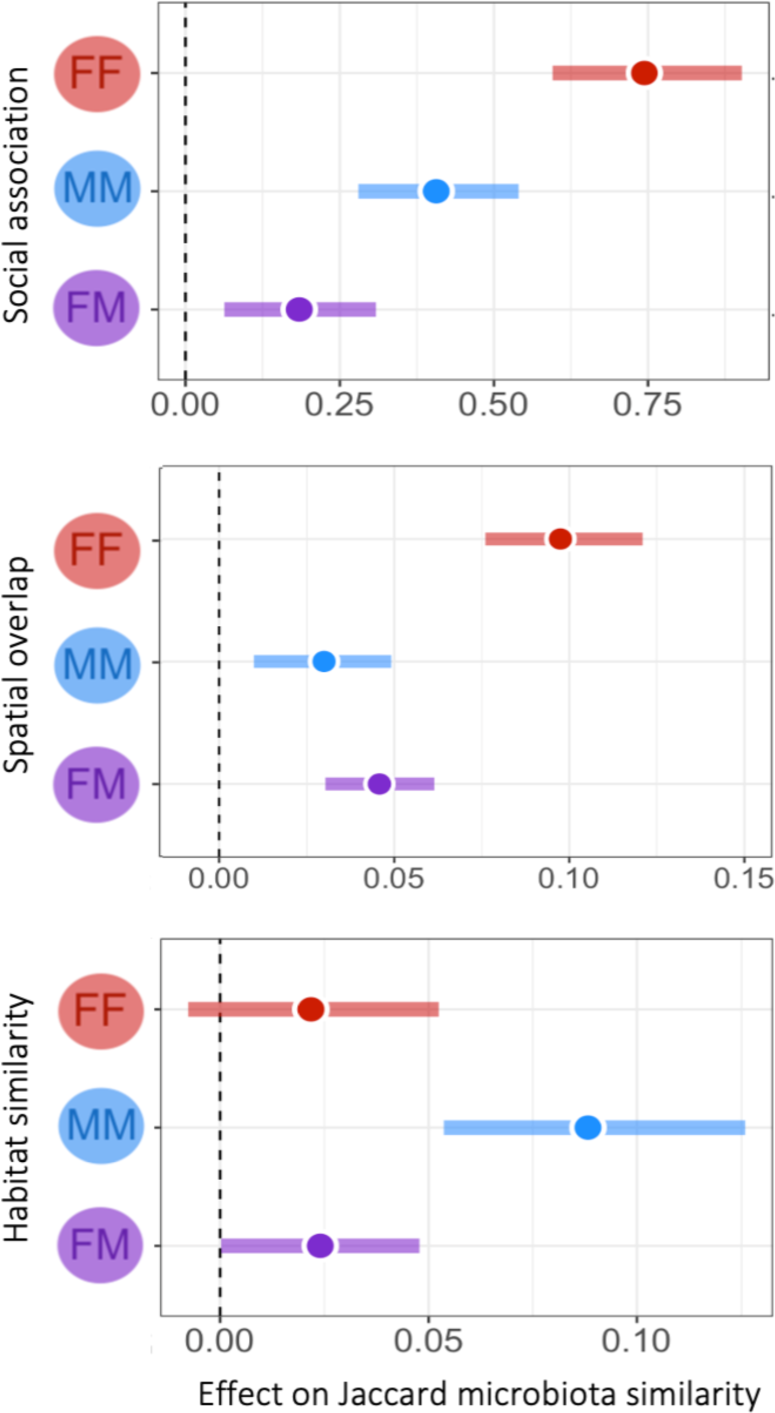
Social, spatial and habitat effects on microbiota across sex combinations. Effects on Jaccard microbiota similarity (x-axis) in pairs with different sex-combinations (colours: FF=female-female, MM=male-male, FM=female-male). Posterior means (points) and their 95% credible intervals (coloured lines) are plotted from Bayesian regression (brms) models (Supplementary Table S2). Where credible intervals do not overlap zero, a social association significantly predicts microbiota similarity.

### Microbial phenotypes predict influence on transmission signals

We used Bergey’s Manual of Systematics of Archaea and Bacteria (54) to classify the aerotolerance and sporulation phenotypes of bacterial genera detected in faecal samples. Since we found that home range overlap predicted gut microbe sharing among mice (Figure 2), we first examined which types of bacteria are detectable in both the local environment and the gut, and thus have the potential for transmission between these two environments. To do this we profiled the soil microbiota from 25 sites across our study area using the same methods used to characterise the gut microbiota, and classified aerotolerance and spore-formation ability for bacterial genera present in the soil. Soil microbiota was more diverse than mouse gut microbiota with 3450 ASVs and 502 genera found from soil compared to 1289 ASVs and 188 genera in the mouse gut (Supplementary Figure S2). We searched for phenotype information for all genera present in mice or in both mice and soil. Of taxa present only in the soil we searched phenotype information for genera found from at least 50% of the soil samples. We found that while few taxa overall were shared between mouse faeces and local soil, with just 6% ASVs and 24% of genera detected in faecal samples also found in local soil, nearly all shared taxa were aerotolerant (Supplementary Figure S6).

Overall, we could reliably infer phenotypic information for 60% of gut microbial genera (Supplementary Table S4). Using this subset, we then used two analytical approaches to assess which kind of bacteria spread through each transmission route in this population. First, we repeated our probabilistic models using Jaccard indices calculated from one of four phenotypic subsets: (i) strict anaerobes (ii) aerotolerant (iii) spore-forming and (iv) non-spore-forming taxa. As these subsets contain varying numbers of ASVs and differ in the mean proportion shared among hosts, we cannot directly compare the strength of a specific effect across models. However, we can assess how the relative strength and credibility of key effects within each model varies according to the subset of microbes being considered. When considering only aerotolerant taxa, the social network’s effect on ASV sharing became weaker and less certain compared to the effects of spatial overlap and habitat similarity, to the extent it was no longer significant (Figure 4, Supplementary Table S5). In contrast, when only strictly anaerobic taxa were considered, the relative magnitude of effects mirrored that observed for all taxa, with the social network predicting sharing of ASVs among mice more strongly than shared use of space or habitat similarity (Figure 4; Supplementary Table S5). Overall, these results suggest that anaerobic taxa predominantly drive the effect of social association observed. In contrast, subsetting microbes by their ability to form spores did not appreciably alter the relative magnitude of any of the effects (Supplementary Table S5, Supplementary Figure S7).

**Figure 4.**
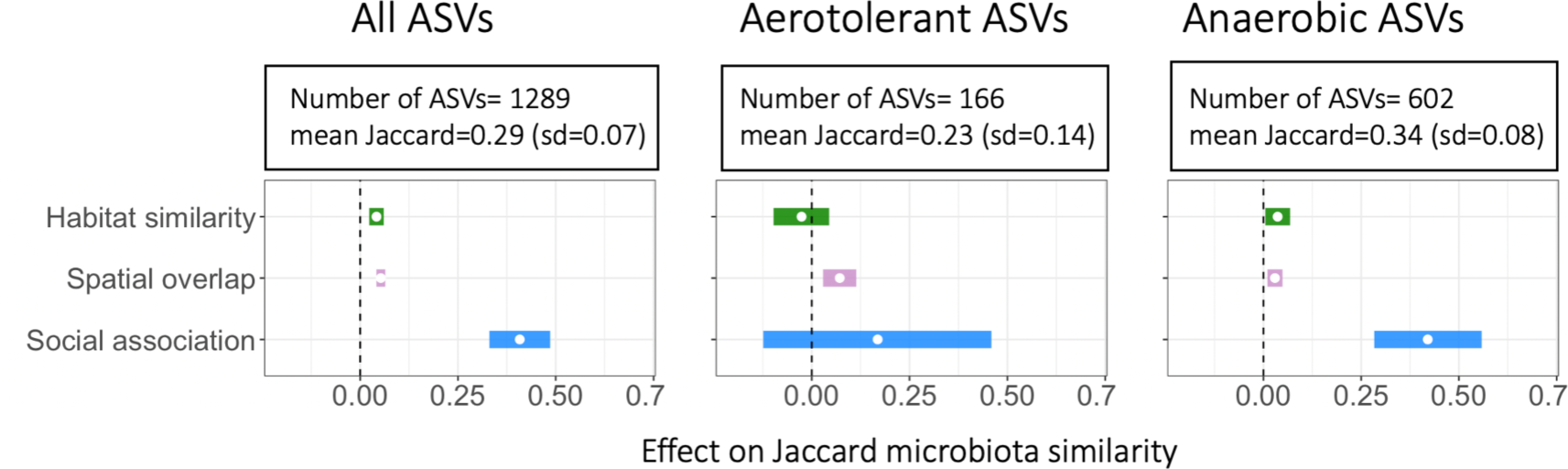
Effects of social association, spatial overlap and habitat similarity on the sharing of either anaerobic or aerotolerant gut microbial taxa. Posterior means (points) and their 95% credible intervals (coloured lines) are plotted from Bayesian regression (brms) models (Supplementary Table S1A, S5A-B) with pairwise the degree of microbial ASV sharing among hosts (Jaccard Index) as the response. Where credible intervals do not overlap zero, a variable significantly predicts microbiota similarity while controlling for other predictors and covariates.

Second, for each bacterial genus we calculated “*importance scores*”, which capture the impact of their inclusion on the model’s ability to detect social, spatial and habitat effects respectively. We did this by dropping each genus in turn from the data, re-calculating the Jaccard index, re-running our main model and measuring the extent to which uncertainty (credible interval width) increased around each effect. ’Important’ genera for a given effect are therefore those which increase the signal-to-noise ratio. In all models where a single genus was excluded, we found social, spatial and habitat effects that were significant and similar in magnitude to those detected in the full model. This indicates that no single genus drove these effects, but rather each genus has a small contributory effect that varies in magnitude and direction. For all three effects (social, spatial, and habitat) importance scores showed no strong phylogenetic clustering, suggesting taxa from across the bacterial phylogeny contribute to each (Figure 5). However, spatial and habitat importance scores were significantly positively correlated (*r*=0.35, p<0.001), suggesting some overlap in the taxa that are most important in generating these two effects (Figure 6A). By contrast, neither spatial nor habitat importance scores correlated with social importance scores, implying different set of taxa were influenced by social vs environmental associations among mice (Figure 6A). Indeed, most of the ten most socially important genera belonged to the phylum Firmicutes and none were from Proteobacteria, while most of the ten most important genera for both spatial and habitat signals belonged to Proteobacteria (Supplementary Figure S8).

**Figure 5.**
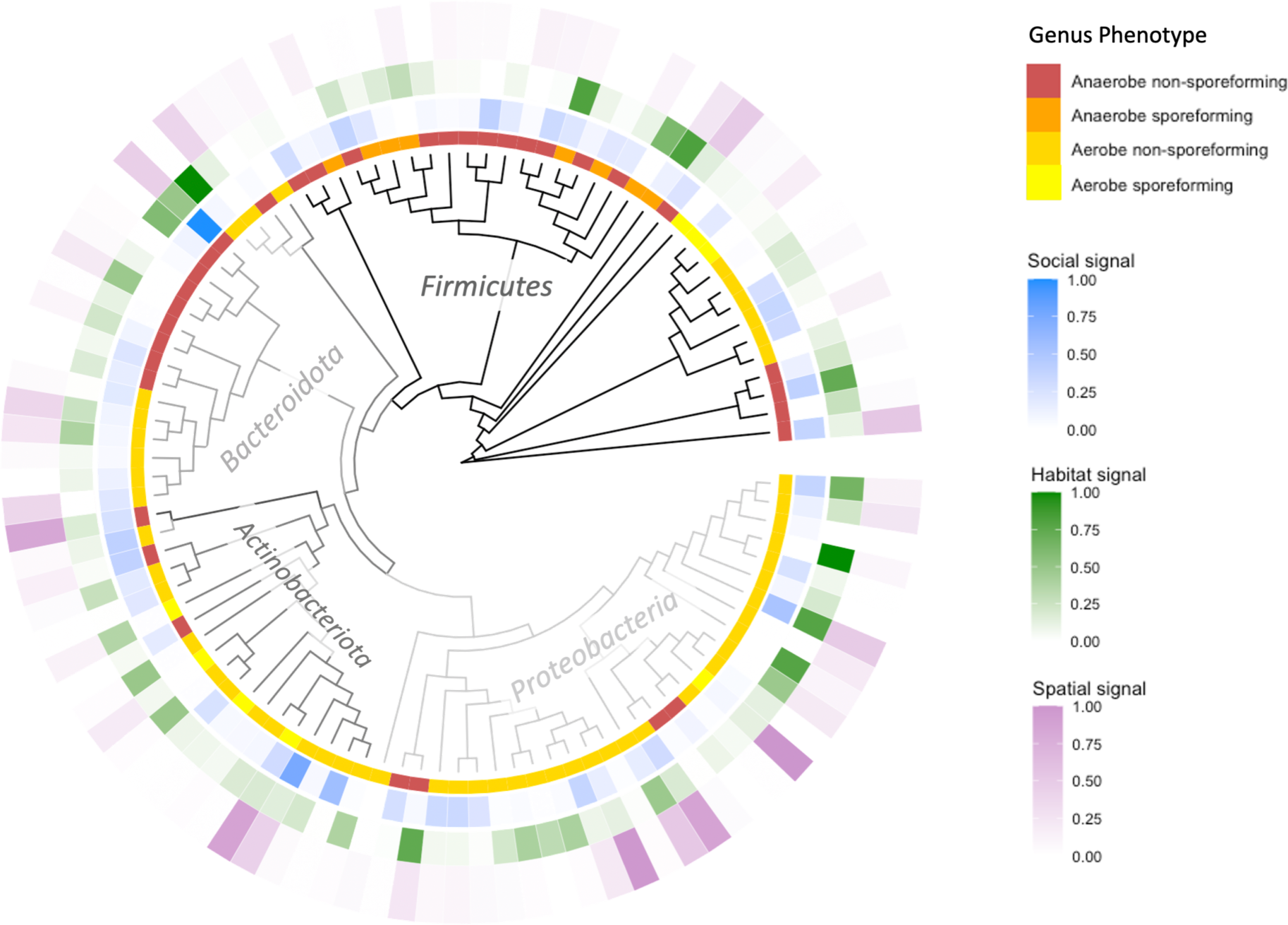
Phylogenetic distribution of bacterial phenotypes and importance values for transmission signals across gut microbial genera. Figure is covering the genera that were included in the models, i.e., the 110 genera with complete phenotype information available out of 188 found in the mouse gut.

**Figure 6.**
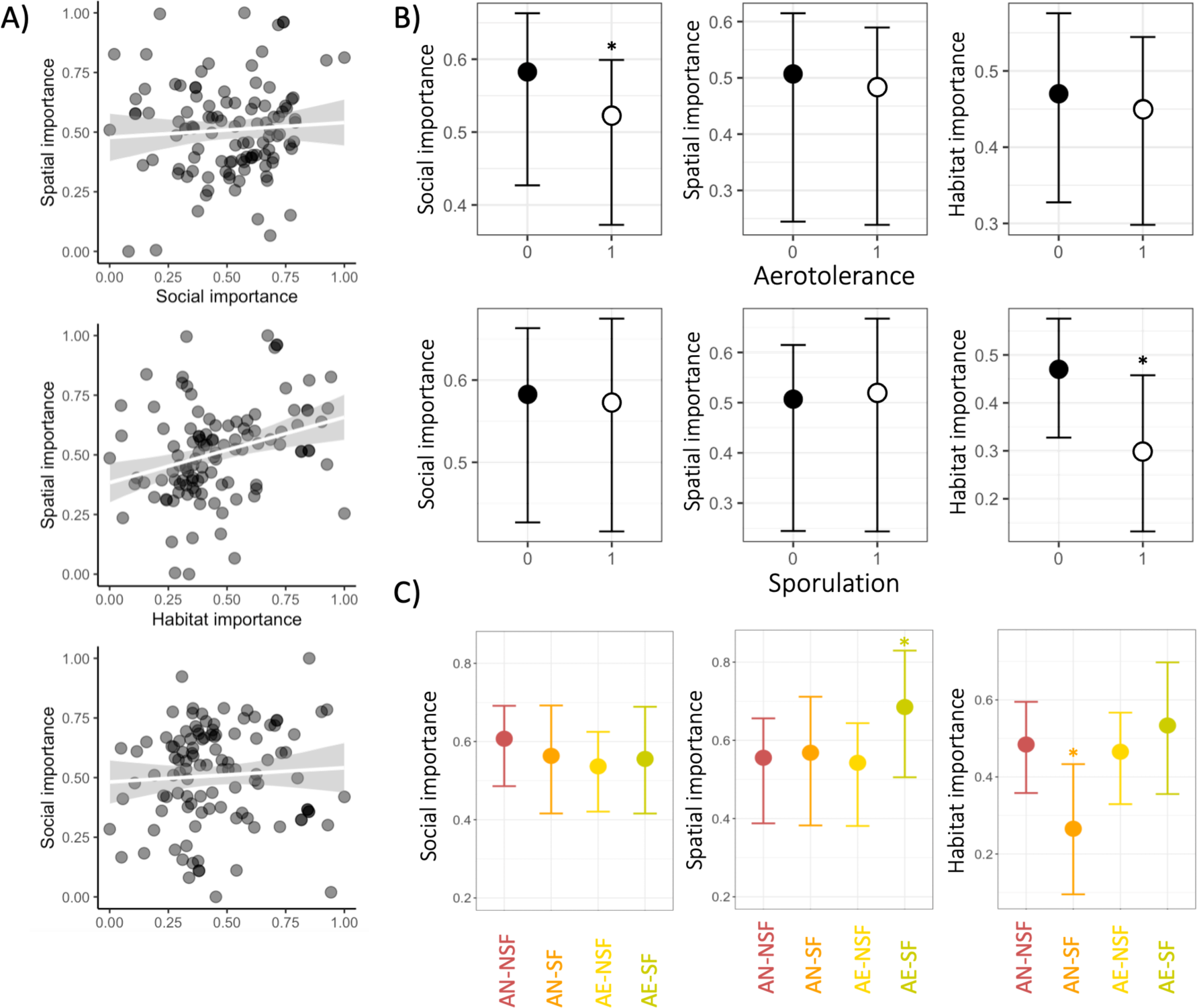
Distribution of importance scores and bacterial phenotypes. A) Correlations among importance scores of social, spatial and habitat signal. B) Statistical Effects of aerotolerance on social, spatial and habitat importance values (y-axis), based on a phylogenetically controlled Bayesian regression model (Supplementary Table S6), predicting importance scores with aerotolerance, sporulation and their interaction across bacterial genera. C) Statistical Effects of phenotype combinations (colours, AE-SF= Aerotolerant spore-former, AE-NSF= Aerotolerant non-spore-former, AN-SF= Anaerobic spore-former, AN-NSF= Anaerobic non-spore-former) on social, spatial and habitat importance values (y-axis), based on a phylogenetically controlled Bayesian regression model (Supplementary Table S7), predicting importance scores with phenotype combination categories across bacterial genera. Phenotypes marked with an asterix differ significantly from other taxa in their values of given importance (Post hoc model, Supplementary Table S8).

We next used Bayesian generalised linear models to formally test whether aerotolerance or spore-forming ability predict the importance of bacterial genera for social, spatial and habitat signals, while controlling for any phylogenetic structure. This revealed that, consistent with the earlier modelling approach, social importance was negatively associated with aerotolerance, (Posterior mean: -0.07, CI from -0.14 to -0.00; Supplementary Table S6A, Figure 6B). Habitat importance was also negatively predicted by sporulation (Posterior mean: -0.17, CI from -0.31 to -0.02). To further explore effects of phenotype interactions, we then ran additional models predicting importance scores with a more detailed combination phenotype as a 4-level factor (aerotolerant spore-formers, aerotolerant non-spore-formers, anaerobic spore-formers, anaerobic non-spore-formers), and post hoc models to determine whether the most important phenotypes had significantly different importance compared to other genera. This revealed that spatial importance scores were in fact significantly higher in aerotolerant spore-formers than other genera (Supplementary Tables S7B, S8B, Figure 6C), while controlling for bacterial phylogenetic relatedness. Social importance was on average highest in anaerobic non-spore-formers but this effect was not significant (Supplementary Tables S7A, S8A Figure 6C). Furthermore, compared to other phenotypes, anaerobic spore-formers had significantly lower importance for the habitat signal (Supplementary Tables S7C, S8C, Figure 6C).

## Discussion

Recent research has shown that mammalian gut microbiota can be influenced by transmission through social behaviours (19) or environmental contact (55), but the relative contributions of social vs environmental transmission pathways have not been explored simultaneously. Here we find evidence for parallel and distinct effects of environmental and social contact transmission in shaping the gut microbiota composition of wild mice. The microbiota of wood mice was affected by both their shared use of space and by social associations with other conspecifics, with these transmission pathways generating effects that differed both in magnitude and in the microbial taxa involved. Specifically, the social signal in wood mouse gut microbiota was over eight times stronger than effects of either spatial overlap or habitat similarity, meaning that mice who were often observed together (in the same place close in time) shared many more gut microbial taxa than those who only shared living space or were exposed to similar microhabitats. Social contacts have been found to homogenise the microbiotas of interacting individuals of many social species (25–28), but the fact that social transmission can have such a strong effect independent of shared space and even in a relatively non-social species such as the wood mouse, is striking. These results concord well with earlier findings on microbiota transmission from another wood mouse population, where with more sparse behavioural monitoring (using only 9 loggers across a similar sized area) we showed that social networks strongly predicted gut microbiota composition, independent of spatial proximity (a simple distance between each mouse’s average point location) (35). Notably, unlike in that earlier study, the findings here are based on spatiotemporal co-occurrence data collected with completely unbaited RFID loggers recording completely natural space use behaviour, suggesting that strong social effects on the gut microbiota are general across wood mouse populations and readily detectable using different tracking methods. The finding that home range overlap and habitat similarity had generally small effects on microbiota sharing is also in line with earlier findings from other wood mouse populations, suggesting that geographic locations 1-6 miles apart explained a relatively small amount of microbiota variation (56,57).

These social and spatial effects we believe reflect transmission processes, while the effect of habitat similarity may reflect a mixture of transmission and selective forces imposed by the environment on microbiota (such as effects of diet or stress on microbiota). Since transmission processes are expected to affect which taxa colonise a host but not necessarily how well they subsequently grow, we expected social and spatial signal to be more readily captured using binary measures of microbiota sharing whereas habitat signal might be captured well by abundance-weighted measure of microbiota similarity (Bray-Curtis similarity). In line with this hypothesis, social and spatial signals were doubly as uncertain in model predicting abundance-weighted Bray-Curtis similarity compared to the model predicting Jaccard microbiota similarity. While habitat signal was also less uncertain, this difference was less striking.

When gut microbes are transmitted through social contact, their distribution across the host population can reflect patterns of social behaviour among hosts. Consistent with this, we found that the social network’s effect on microbiota composition varied between sexes and across seasons. The social effect on microbiota composition was strongest for female-female pairs, weakest in female-male pairs with male-male pairs having an intermediate effect size. The social effect was particularly strong in female-female pairs during spring (Feb-Jun), when social associations were stronger on average. This seasonal difference in the effect of social association on the microbiota may be linked to behavioural differences of wood mice across their breeding cycle. During the breeding season (approx. June-November in Wytham), wood mice, especially females, are more solitary and territorial compared to the non-breeding season, when multiple mice may co-nest together in same-sex groups (58–61). Interestingly, the pattern of sex-dependency in the social transmission effect detected here differed from earlier findings from another wood mouse population, where social association predicted microbiota similarity only in male-male and male-female, but not female-female pairs (35). While more detailed behavioural data would be needed to understand which specific social behaviours are involved in gut microbial transmission, these findings already emphasize that differences in the ecological and social environment among populations of the same species can change the way individuals exchange microbes, as microbial transmission may depend on fine-scale variation in social relationships, even among individuals utilizing the same space.

Our results clearly showed that not all microbes are equally spread by different transmission routes. Through two complimentary analytical approaches, we found that different microbial taxa underpinned social effects on the microbiota compared to effects of spatial overlap and habitat similarity. This means that different members of the gut microbial community may be transmitted through social contacts and a shared environment respectively. Indeed, we found the social transmission signal was specifically driven by sharing of anaerobic bacteria, as it was no longer significant when they were removed from the analysis, whereas the spatial transmission signal was detectable among both aerotolerant and anaerobic taxa, and if anything was slightly stronger for aerotolerant bacteria. Deconstructing the whole-community-level effects into genus-level contributions further revealed that spatial signal in the microbiota was driven most strongly by aerotolerant spore-forming genera, while the social signal was most influenced by anaerobic genera, most strongly anaerobic non-spore-formers. Sampling the microbiota of the environment (soil), also revealed that the microbes present in both soil and mouse gut were rather exclusively aerobes. We do not expect these soil microbes to represent a full picture of microbes present in the mouse environment, but the fact that only aerobic bacteria seemed to spread between soil and mice supports the idea that aerotolerance is important for environmental transmission of gut microbes and that anaerobic taxa may need a different mode of transmission. In future, more thorough sampling and source-tracking approaches could identify through which substrates indirect host-to-host transmission happens. Similar evidence supporting either social transmission spreading less oxygen-tolerant taxa, or environmental exposure spreading more oxygen-tolerant bacteria exists in the literature. Studies of wild baboons, for instance, found that social associations based on the grooming social network predicted microbiota similarity and this effect was driven by anaerobic and non-spore forming bacteria (28). A follow-up study further showed that baboon populations living in different geographic locations differed specifically in the aerobic microbiota they hosted (47). Studies on the human gut microbiota have also reported findings that suggest links between bacterial phenotypes and transmission ecology. For example, gut microbial taxa which can form spores are more prevalent among people than those which cannot, consistent with them more readily transmitting among hosts (50) and microbial strains shared among people from geographically distant locations (i.e., between populations) were more likely to be aerotolerant and spore-forming than those shared among household members (21). Alternative evidence comes from a laboratory rodent experiment, where aerotolerance was shown to be associated with transmission pathways of gut microbes among 17 inbred laboratory mouse lines derived from geographically distinct wild populations (62). Here, microbial transmission between adult mice (“horizontal transmission”) was driven by aerotolerant taxa, while obligate anaerobes were found to be only vertically transmitted (passed from mothers to offspring in birth). However, in this study, horizontal transmission of microbes was mostly limited to transmission through the aerobic environment, as caged mice did not socially interact with each other. Future research could usefully assess whether vertically transmitted gut bacteria overlap with those taxa transmitted horizontally through intimate social contacts later in life. If so, this would mean that the same (perhaps anaerobic, non-spore-forming) microbial taxa spread through both social contacts and from mother to offspring. Over evolutionary timescales, this kind of transmission ecology might lead to anaerobic and/or non-spore-forming microbes getting stuck not just inside a host social network but also the branching tree of host lineages, such that those microbes which are transmitted through the most intimate interactions become most specialised in their host species. Supporting this, the most host-specific gut microbial taxa were recently found to be enriched in anaerobic phenotypes across mammalian hosts (63). A recent simulation study also suggested that gut microbial transmission mode (horizontal vs vertical) could predict the level at which the bacteria may establish a stable host-microbe relationship across evolutionary time, with microbes less able to persist outside the host evolving more host-specialist life-style (64). However, like the above-mentioned laboratory mouse study (52), this simulation study contrasted maternal “vertical transmission” with general “horizontal transmission”, pooling together all microbial transmission processes among adult animals, thus not considering the separate effects of social and environmental pathways. Rather than contrasting vertical and horizontal transmission, future research might benefit from categorising transmission processes into maternal, social and environmental pathways. This way we can separate environmental transmission that may spread more generalist gut microbes from more intimate transmission routes (maternal and social) which may spread more host-specialised gut microbes. Consistent with the idea that maternal and social transmission may spread similar bacteria, our recent study on wood mice found that as young mice age, maternal transmission processes are gradually replaced by social transmission processes (65), with maternal and social effect in these mice both driven by microbes in the same family Muribaculaceae (35,65).

If aerobic and spore-forming vs. anaerobic and non-spore-forming microbes spread from host to host through somewhat different transmission pathways as our data suggest, this has two important implications. First, anaerobic non-spore-forming microbes that require more intimate transmission routes may be more likely to evolve a more stable relationship with their host (as suggested by Leftwich et al., (64)) and perhaps more mutualistic relationship as well, since they are more dependent on their host species (as suggested by Moeller et al., (62) based on Brown et al., (66)). Compared to environmental transmission, social transmission may therefore be expected to spread microbes with greater functional significance for the host, for example in nutrition (67) or protection against pathogenic infection, as has been shown in some insect systems (68). Social transmission of key symbionts is an important possibility to consider when weighing the pros and cons of social lifestyle in theory concerning the evolution of sociality, especially if socially acquired microbes are among those that influence social behaviour (7), as this could potentially lead to positive feedback loops in which social behaviours spread social behaviour-boosting microbes (6). Second, the fact that some microbes only live in and transmit between hosts while others readily spread between the host and the external environment calls in question the relevance of viewing host associated microbiotas as classic metacommunities, like island ecosystems. While some assumptions in classic metacommunity ecology (such as the idea of completely inhabitable matrix separating habitat patches) have been updated to better model the microbiota (46), assuming that all members of a microbiota can similarly persist in the environment between hosts seems unrealistic. Microbial taxa may vary greatly in the extent to which they experience the host as a true island. For instance, anaerobic microbes may well live in a strict metacommunity, structured by the host social network, while the aerotolerant microbes may experience a much more continuous landscape, more analogous to valleys amidst hills than islands in the sea. Similar variation among species in how they experience a metacommunity landscape of course also exists in macroecological metacommunities, for example due to varying dispersal abilities (69). However, in microbiotas a key difference may be that this variation arises not only from varying abilities to cross the matrix but rather a gradient of abilities to persist and live within it.

Going forward, to better understand the transmission ecology of different members of microbiota will require further work to characterise microbial phenotypes associated with the ability to persist outside the host. Data on aerotolerance and spore-formation is still lacking from many gut microbial genera of wild animals. For example, the most common genus in our dataset was an unnamed group of Muribaculaceae, a family which has itself only recently been named and characterised (70). In reality, this group may consist of multiple different genera, but with current taxonomic profiling tools, it is impossible to delineate them. Furthermore, aerotolerance and spore-formation are probably not the only relevant phenotypes determining gut microbial transmission pathways. For instance, persistent states mediated by toxin-antitoxin system, metabolically sparing “viable but non-culturable” (VBNC) states, or morphological adaptations in the cell wall could all potentially affect the environmental transmissibility ability of gut microbes (50). In fact, a recent thorough exploration of the human gut microbiota transmission landscape found that cell wall properties (as described by gram stain) were associated with human-to-human transmissibility of gut microbes (21). As culture-based phenotypic information can be limited, especially in wild host species, there is also a lot of promise in the growing number of tools developed for predicting bacterial phenotypes from genomic data (See for example TRAITAR (71)). Here, a potentially useful but so far unexplored method for classifying aerotolerance phenotypes from bacterial sequence data could involve characterising Ribonucleotide Reductase enzyme genes (72).

Based on our findings, it seems that while gut microbes can be shared among hosts via multiple independent routes (social interactions, and shared space use), the subset of using each of these transmission pathways differ both taxonomically and in traits that affect their ability to persist outside a host. This highlights the need for further research on the transmission dynamics of not only pathogens but also commensal members of the microbiota. Humans are a socially flexible species capable of large-scale modification of their own social contact network, as evidenced by widespread social isolation practises implemented during the SARS-CoV-2 pandemic. Reducing social contact is an effective way to reduce pathogen spread, but multiple studies have also highlighted that we know essentially nothing about the consequences of social isolation for our commensal microbiota and microbiota-mediated health (19,73,74). At the same time, a growing body of evidence emphasizes how diminishing contacts with the natural environment among urbanised human populations can have negative health consequences through a lack of natural microbiota transmission from biotic environments and consequent disruptions to immune development (36–39,75). If isolation from natural sources of environmental microbiota transmission can compromise host health through immune disruptions, what might be the consequence of isolating from natural sources of social microbiota transmission? More research is needed to answer this important question. Notably, both reduced natural contact and reduced social contact are common aspects of modern lifestyles. Such isolation from social and ecological environment may independently disrupt the co-evolved transmission networks of human microbiota (74), reflected by the observation that modern lifestyles are linked with a range of immune disorders (76,77)and seem to be depleting the diversity of human gut microbiota (78,79).

## Methods

### Field data collection

During July-November 2019, we collected faecal samples and tracked the movements of 164 wild wood mice living within a woodland plot (Holly Hill) in Wytham Woods, Oxford, UK (51.77 °N, -1.33°S). This involved fortnightly trapping to tag mice and collect samples, alongside continuous passive tracking of tagged individuals using RFID technology. The focus study area, where mouse behavior was tracked, was a 2.56 ha (160m x 160m) “core grid”, but to minimise edge effects mice were trapped and tagged from an area larger than this core grid, from a 4 ha (200m x 200m) “extended grid” spanning up to 40 meters outside of the core. Trapping sessions were carried out in the area from November 2018 to November 2019, with captured mice aged and sexed, and injected with a subcutaneous PIT-tag for permanent identification and tracking. After processing, all individuals were immediately released at the exact location they were trapped. Faecal samples for microbiota analysis were collected from the traps of identified individuals into sterile sample tubes with sterile tweezers and frozen at -80°C within 4 hours of collection. All traps showing signs of rodent presence were carefully washed and sterilised in bleach solution before the next trapping session, to eliminate cross-contamination. Additionally, in the beginning of the study period (between November 2018-February 2019), 25 soil samples were collected from around the 4 ha extended grid to serve as a general reference to the local soil microbiota. A soil sample was collected by digging a spoonful of soil (∼200 mg) from 3 cm underground, creating a mix from three digging spots within a meter of a mouse trapping location.

Mouse behaviour was monitored with a set of 60 custom-built RFID-loggers distributed across the study site, recording the time-stamped presence of any individual that came within its read range (∼1m^2^). Loggers were unbaited, positioned evenly across the grid and rotated fortnightly to ensure even spatial coverage of the area. This scheme meant that each 10 x 10 m grid cell of the study site was covered by a logger for a fortnight every two months, i.e., 25% of the time. In addition to these evenly spaced “above-ground” loggers used to derive social networks, to derive home range estimates we also included data from an extra set of 60 loggers positioned at entrances to mouse burrows between July-November. These “burrow loggers” were not rotated but were distributed approximately evenly across the study area. Further details of logger devices and the tracking protocol can be found in Supplementary Figure S9.

Shortly after the study (May, 2020), we completed a thorough survey of vegetation and microhabitat variation across the study site, in which the percentage cover by each of the eight main ground cover types in the area was recorded for each 10 x 10m grid cell of the plot (Supplementary Figure S10).

### Social network construction

Social networks were constructed using data from all 60 above-ground loggers for the full 10-month period (‘Full Social Network’) as well as separately for spring (Feb-Jun) and Fall (Jul-Nov) (‘seasonal networks’). This division of the year into “seasons” was done based on cutting the whole study period in two equal halves, but it also approximately mirrors the natural seasons of wood mouse breeding; wood mice are reproductively more inactive during late fall/early spring and become increasingly reproductively active during late spring and breed until early fall. Consistent with this, juveniles started appearing in the data only in early September and stopped appearing in turn of November-December, meaning that females were typically pregnant from August onwards. While reproductively active, female wood mice are less social and more territorial (60,61,80). Networks were constructed and visualized using our custom social network inference and plotting functions in R (see github: https://github.com/nuorenarra/Social-Network-Analysis) with the help of R package *igraph* (81). These functions took the logger data, consisting of time-stamped observations of tagged individuals in fixed locations, and calculated a pairwise association index for all mouse pairs based on the frequency with which they were observed at the same location during the same short time window. Logger data was first filtered to include only the normal hours of activity for this nocturnal species (16.00-08.00). Each of these 16hr logging ‘nights’ formed the primary units for social association inference. For each night, a pair of mice were considered ‘associated’ if they were observed at the same location within 12 hours of each other, consistent with our previous work (Raulo et al. 2021). These instances of association were then used to calculate an association index defined as:

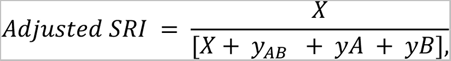

where *X* = the number of nights in which individuals *A* and *B* were observed associated (in the same location within 12 h of each other), *y_AB_* is the number of nights in which *A* and *B* were *both* observed but not associated (observed at the same location but more than 12h apart), *y_A_* and *y_B_* are the number of nights in which both were known to be alive but only *A* or *B* was observed respectively. Accounting for lifespan overlap in this metric allows us to more accurately summarize the temporally fluctuating social structure of the mouse population in one static social network.

### Estimating home range overlap

An animals’ home range can be defined as “the area, usually around a home site, over which the animal normally travels in search of food” (Burt, 1943) and is commonly presented as a utilization distribution describing the probability of space use with respect to time (Powell & Mitchell, 2012). We quantified home ranges from logger data using an autocorrelated kernel density estimator (“AKDE”; (Fleming et al., 2015) implemented using the *ctmm* package (Calabrese et al., 2016). Home range boundaries were delineated at the 75% level to provide an estimate of the core home range, the smallest area that one could expect to find a given individual inside, with 75% probability. In other words, each individual mouse’s home range was described as a three-dimensional probability distribution of space utilization, where the two base dimensions were actual space and the third dimension was utilization intensity, i.e. how frequently the mouse used a given region within its range. Home ranges were calculated only for individuals satisfying our criteria for a complete and stable observation record, based on variograms estimating temporal autocorrelation in spatial records (Supplementary Appendix S1). Under these criteria, home ranges could be estimated for 104 of the 157 mice recorded on loggers. Among these 104, we calculated home range overlap for each mouse pair using the “overlap” function in *ctmm* and the Bhattacaryya coefficient, defined as:

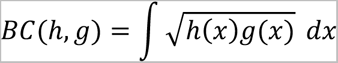

, where *h* and *g* are two probability distributions of value *x* respectively.

Since reliable home range estimation requires a considerable amount of tracking data, home ranges were estimated using all available logger data from both above-ground and burrow loggers. To ensure that higher logger density in the fall (when burrow loggers were deployed) did not bias home range estimates, we used a subset of well sampled mice from the fall to show that home ranges calculated from the sparser above-ground logger data were overlapping and comparable in size to home ranges based on all logger data pooled for the same individuals (see Supplementary appendix S1).

### Estimating habitat similarity

Habitat similarity between mice was estimated using data on the percentage cover by each of the eight main ground cover types within each mouse’s home range (75% core kernel density area). The main ground cover types were defined as: 1) open ground (OG; no plant coverage), 2) dog’s mercury (DM; covered by *Mercurialis perennis*), 2) bluebell (BB; covered by *Hyacinthoides non-scripta*), 3) bramble (BR; covered by *Rubus fruticosus*),4) grass (G; covered by grass species in family *Poaceae*), 5) sedge (S, covered by *Carex pendula*), 6) Enchanter’s night shade (EN; covered by *Circaea lutetiana*), 7) wild garlic (WG; covered in *Allium ursinum*) and 8) Currant (RI, covered by *Ribes spicatum*) (see Supplementary Figure S10). For each mouse, we calculated normalized abundance for each ground cover type, as the sum of its coverage across the home range (in m^2^) divided by home range area. Using these values, we then used package *vegan* to calculate habitat similarity for all pairs of mice, using the Bray Curtis Index (82).

### Microbiota profiling

We profiled microbial communities by extracting DNA from faecal and soil samples and using primers 515F and 926R (83) to amplify and sequence the V4-V5 region of the 16S rRNA gene in bacteria/archaea. Full details of the laboratory work, library preparation and sequence data bioinformatics can be found in Supplementary appendix S2. In brief, we used the DADA2 algorithm to infer microbial sequence variants (ASV) from the sequence data and assigned taxonomy using the SILVA database (version 138) after which the data was decontaminated (84) and filtered to remove non-gut-microbial taxa and samples with low read counts. Finally, abundance data was normalized to proportions of each ASV per sample.

## Statistical analyses

### Describing microbiota variation

To characterise variation in microbiota composition among individuals (beta diversity), we used the Jaccard index, which captures the proportion of microbial ASVs detected across a pair of individuals, that are shared between them. This metric provides an intuitive way to capture transmission signals (as transmission should affect the presence/absence, but not necessarily the relative abundance of taxa within a host. Previous work suggested that abundance-weighted measures of beta diversity (e.g. Bray-Curtis dissimilarity) do not provide better detection of microbiota transmission signals compared to the Jaccard Index (35). To ensure Jaccard index was the most optimal beta diversity metric for picking up transmission signals in the data, we also modelled alternative measures of microbiota sharing: abundance weighted Bray-Curtis microbiota similarity and the raw count of shared ASVs between a pair (scaled between 0 and 1).

To estimate the amount of gut microbiota variation accounted for by stable differences between host individuals vs. temporal fluctuations within host individuals, we used Principal Coordinates Analysis (PCoA) and marginal PERMANOVAs (implemented with the adonis2 function of package *vegan* (85) to predict the Jaccard Index across samples from repeatedly sampled individuals (n=255 samples from 82 individuals with a mean 3.1 samples per individual, range 2-10), using host ID and sampling month (as a factor) as fixed effects.

### Modelling the effect of different transmission pathways on microbiota

To test the effects of social and environmental transmission on microbiota composition, we constructed a model in which microbiota sharing among pairs of mice is predicted by their social association strength, spatial (home range) overlap, and habitat similarity. Here, the effect of social association while controlling for the effects of spatial overlap and habitat similarity is meant to capture the effect of social contact transmission on microbiota. Similarly, the effect of spatial overlap controlled against the other two main predictors is meant to capture the effect of microbial transmission from and through shared space, and the effect of habitat similarity controlled against the other two predictors is meant to capture the effect of convergent exposure to similar environmental pools of microbes (those predicted by similar vegetation) and similar selective forces shaping microbiota (e.g., diet). To build this model we used a dyadic Bayesian multilevel model framework implemented in package *brms* (86), as validated and described in (35) (See also https://github.com/nuorenarra/Analysing-dyadic-data-with-brms). This model framework allows “multi-membership” random effect structures that can account for the types of dependence inherent to pairwise comparisons as well as repeated sampling of the same individuals (87). Models used dyadic measures of microbiota similarity (Jaccard index, Bray-Curtis similarity or number of shared taxa) as the response variable, including all sample pairs except those from the same individual mouse. These values were modelled as a function of the predictor variables described above together with a set of technical and biological covariates: host age class similarity (same vs different), sex similarity (same vs different), time interval in days between samples, sample extraction distance (the physical distance between two samples on plates during DNA-extraction, as described in Supplementary Appendix S2), read depth difference and PCR plate similarity). Models with Jaccard index of Bray-Curtis as the response used beta regression (likelihood family = Beta) where the response is a proportion.

Where number of shared taxa was the response, a poisson regression was used (likelihood family = Poisson). The models included a multi-membership random effect (random intercept) for the identity of samples (Sample A + Sample B) as well as individuals (Individual A + Individual B) involved in each pairwise comparison. In addition to this primary model, we ran one additional pair of models to explore seasonal and sex-specific differences in social and environmental influences on the microbiota. Here, we modelled Jaccard Index as a function of the same set of predictors but for each of the seasonal data subsets of the social network (spring or fall), and included an interaction term between sex-combination (3-level factor: female-female, female-male, male-male) and social association. The brms model uses a Markov chain Monte Carlo sampler (Hamiltonian MC sampler, implemented with RStan (88) a wrapper for Stan, (89) to estimate posterior distributions (87). We ran the models with 4 parallel chains, each with 1000 warm-up samples preceding 4000 actual iterations and used posterior checks to ensure reliable model performance (90). Specifically, we ensured that the chains converged, Rhat values were <1.05, Bulk effective sample sizes were no smaller than 10% total posterior draws, and the sampler took small enough steps (adapt_delta=0.98, max_treedepth=13) to avoid excess (>10) divergent transitions after warm-up.

### Transmission signals and microbial phenotypes

We used Bergey’s Manual of Systematics of Archaea and Bacteria (54) to classify the aerotolerance (aerotolerant or strictly anaerobic) and sporulation (spore-forming or non-spore forming) of each bacterial genus identified. When this information could not be found in Bergey’s Manual (for example, as for some newly named taxa), we sought it from original research papers describing the genus or specifically assessing aerotolerance or sporulation of bacterial genera. Full phenotypic trait data for each genus and data sources are presented in Supplementary Table S1. In our analyses including bacterial phenotypes, we only included ASVs belonging to the 188 genera (out of 234 genera in the full dataset) where aerotolerance and sporulation were both well known. For unknown genera in a given family, we only included in the analyses if there was substantial evidence that all members of that family were of a given phenotype. All genera with unknown, uncertain, or variable phenotypes in terms of either aerotolerance of sporulation were excluded from the analysis. We used this data in two analyses to examine how different transmission signals (social, spatial, and habitat) were related to microbial phenotypes. First, we calculated additional Jaccard indices that reflected the proportion of shared ASVs among those belonging to four phenotypic subsets: (i) strict anaerobes (ii) aerotolerant (iii) spore-forming and (iv) non-spore-forming. These Jaccard Indices were then used as response variables in *brms* models with social, spatial and habitat variables together with covariates, as described above. The different phenotypic subsets of microbiota contained varying numbers of ASVs and differed in their mean similarity across hosts (See Figures 4 & S7). Thus, we cannot directly compare the effects between models predicting these different versions of Jaccard, because they come from data sets with inherently different uncertainty, intercepts and slopes. However, we can assess how the relative strength and certainty of key effects within each model, varies across models. Because these phenotype-specific Jaccard values also included a few zeros (no taxa shared between two samples), to meet beta-regression criteria, Jaccard values were scaled by (Jaccard * (n−1) + 0.5) / n where n is the sample size.

Second, we quantified the importance of each bacterial genus in driving each transmission signal, and then asked whether microbial phenotypes predicted variation among genera in this importance. ‘Importance’ scores for each of the 188 bacterial genera were calculated by dropping each in turn from the microbiota data, recalculating the full Jaccard Index and re-running the above-described brms model (see https://github.com/nuorenarra/Analysing-dyadic-data-with-brms). The “importance score” for each effect of interest (social association, spatial overlap and habitat similarity) reflected the extent to which dropping a genus from the analysis reduced the precision of that effect estimate. Specifically, the Importance of genus *G* for effect *E* was calculated as the increase in the 95% credible interval width *(CIw)* when G is excluded *(CIw_excl_ -CIw_incl_)* relative to the baseline credible interval width when G is included (*CIw_incl_),* divided by the square root of the number of ASVs (*n.ASV*) assigned to genus G:

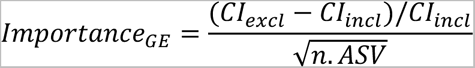

The resulting values were approximately normally distributed, and were scaled between 0 and 1 to create importance scores that were on the same scale as other binary predictors in each model. Across the 188 genera, we tested whether aerotolerance (0/1) or sporulation ability (0/1) predicted importance scores for each effect of interest (social, spatial or habitat), using a *brms* gaussian multilevel model. To control for phylogenetic non-independence among genera in these phenotypes, we also ran the model including the phylogeny among genera (in form of a variance-covariance matrix) as a random structure in the model.

## Funding statement / acknowledgements

This work was supported by a NERC fellowship to SCLK (NE.L011867/1) and funding from the European Research Council (ERC) under the European Union’s Horizon 2020 research and innovation programme (grant agreement n° 851550). AR was supported by a Clarendon Scholarship.

## Data availability

All data used in this study is publicly available through Environmental Information Data Centre (DOI). Microbiome sequence data is additionally located in European Nucleotide Archive (DOI).

