## Supplementary Material for "Social and environmental transmission spread different sets of gut microbes in wild mice"

### Index

#### Supplementary Appendices

**Appendix S1.** Details on home range estimation

**Appendix S2.** Details on gut microbiota analysis

#### Supplementary Figures

**Figure S1.** Examples of social and environmental contact behaviours in wood mice

**Figure S2.** Composition and variation in wood mouse gut microbiota and the local soil microbiota.

**Figure S3.** Variation and co-variation among social association, spatial overlap and habitat similarity.

**Figure S4.** Effects on alternative measures of microbiota similarity.

**Figure S5.** Effects of social association on microbiota similarity across seasons and sexes

**Figure S6.** Phenotype distribution among bacterial genera found in mouse gut, soil or both

**Figure S7.** Effects of main predictors on microbiota similarity in different phenotypic subsets of the microbiota – spore-formation.

**Figure S8.** Overview of the top-ten most important genera for social, spatial and habitat signal in microbiota similarity.

**Figure S9**: The RFID logger device and tracking protocol

**Figure S10**: Main ground cover habitat types in Holly Hill study site.

#### Supplementary Tables

**Table S1.** Model results: Effects of Social association, spatial overlap and habitat similarity on microbiota similarity.

**Table S2.** Model results: Effects of social association, spatial overlap and habitat similarity on microbiota similarity am2ng pairs with different sex categories.

**Table S3.** Model results: Effects of Spring and Fall social association on microbiota similarity among pairs with different sex categories.

Table S4. Aerotolerance and spore-forming ability of bacterial genera together with their importance scores in models where excluded.

**Table S5.** Model results: Effects social association on Jaccard similarity based on aerotolerant vs. anaerobic taxa.

**Table S6.** Model results: Effects of phenotypes on importance scores across bacterial genera.

**Table S7.** Model results: Effects of microbial combination phenotypes on importance scores across bacterial genera.

**Table S8.** Model results: Results of post hoc models testing whether specific phenotype categories differ significantly from other genera in their importance.

#### Appendix S1. Details on home range estimation

For ease, figures mentioned in this appendix are included within this appendix and referred to alphabetically.

Home ranges were inferred from the logger data aggregated within ten-minute periods, such that all records of an individual at the same location within each ten minute-period were considered a single unique "logger visit". To ensure that data quality was suitable for home range analysis we first filtered out all individuals that did not have at least five unique "logger visits" in at least three unique locations. We then inspected the remaining data using empirical variograms, a method visualising the autocorrelation structure in locational timeseries by presenting the semi-variance (proportional to home range size) as a function of increasing time lags, with clear asymptotes diagnostic of range-residency (Fleming et al., 2015, See Figure A). Here, a lack of asymptote indicates either the data is too incomplete to reveal the home range extent, or that the animal is not exhibiting site fidelity (Calabrese et al., 2016). Based on this, we excluded individuals who's variogram showed a positive linear relationship between estimated distance and time lag between logger observations, as these were considered signs of either logger histories that were too incomplete for reliable home range estimation (e.g. due to small amount of data or edge effects on the study grid) or spatially shifting home range location (unstable home range). Under these criteria, home ranges could be estimated for 104 of the 157 mice recorded on loggers.

To estimate home range, for each individual we first fit a series of range-resident movement models using model selection to identify the best model fit, given the data (Fleming et al., 2015) following the workflow described by (Calabrese et al., 2016). We applied the AKDE to these movement models using an error-informed model (with error set to be 1 meter, approximately the detection diameter of loggers). The method accounts for error by first Kriging data to reduce location error before placing the kernels (Fleming et al., 2018; Fleming & Calabrese, 2017). Although the observation model of fixed station data is different to tracking data that the ADKE was developed for, the scales of movement observed for mice was larger than the grid resolution. Accordingly, visual inspection of the kernels against the grid setup and tracking data showed little evidence of excess density at the stations.

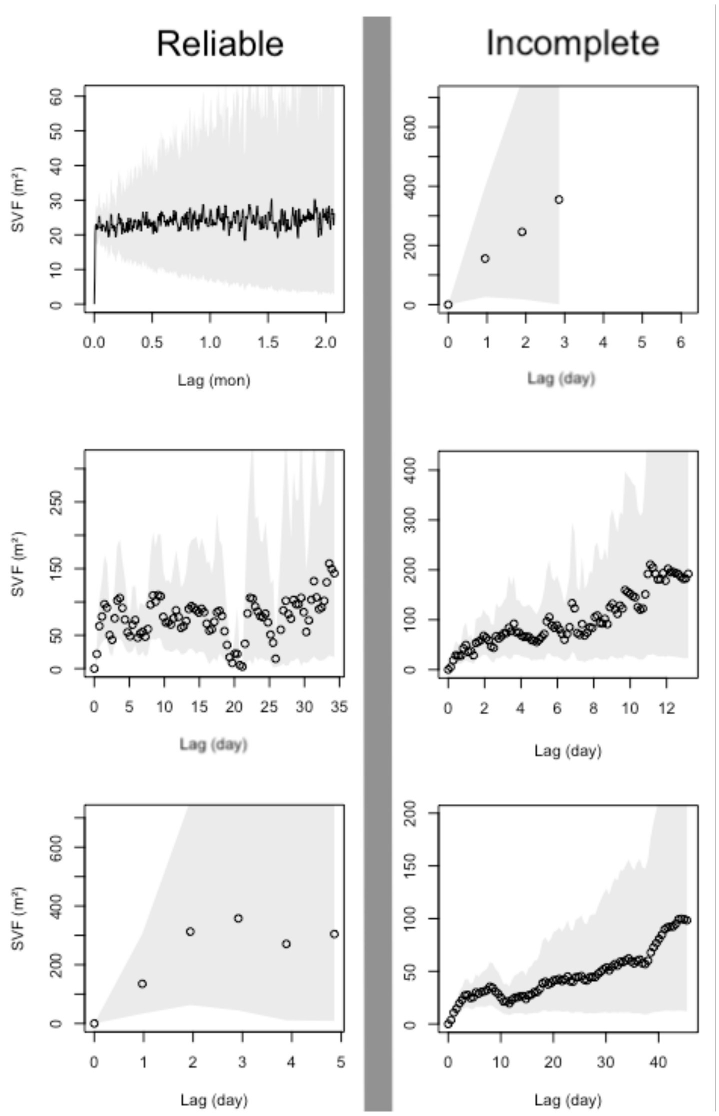


**Appendix S1 – Figure A.** Examples of variograms used as an inclusion criterion for home range estimation. Six example variograms describing the completeness and stability of location-observation record of six individuals. Variograms were used to filter out individuals with insufficient logger data for reliable home range estimation. Here, each box is the observation record of one individual mouse, where x-axis describes time lag in days/months between logger observations and y-axis (SVF) describes the expected distance between observations. Where the distance estimate grows linearly with time-lag between observations (right-hand panel), individual's home range is unstable (shifting) or observation data is not complete enough to capture the true home range and thus estimates for these individuals are not reliable.

*Validation of using logger data with varying density for home range analysis.*

As movement-model based home range analysis requires an extensive number of relocation data points for an individual, we included observations from all logger data for home range analysis to maximize the number of individuals for whom home range could be reliably estimated. This induced a sampling inbalance in our data, as we had more loggers out covering the same area during Fall (n= 60 evenly spaced above-ground loggers, July-November) than in Spring (n= 60 evenly spaced above-ground loggers and 60 burrow-loggers, February-June). To ensure this was not biasing our home range estimates between seasons, we chose a subset of well-tracked mice from the fall period and created a comparable pair of home ranges for each individual using 1) the full logger data (dense data home ranges) and 2) subset of logger data from the 60 above ground loggers only (sparse data home ranges). Across example mice, these two home range estimates were largely overlapping (See Figure B below) and had area estimates that were highly correlated (r=0.97, p<0.001; Figure C below). This implies that our initial criteria for data density acceptable for reliable home range estimation were realistic, estimates were asymptotic and thus including any additional observation data had negligible influence on the estimates. Based on this, we decided to use all available logger data to infer home ranges for as many mice as possible.

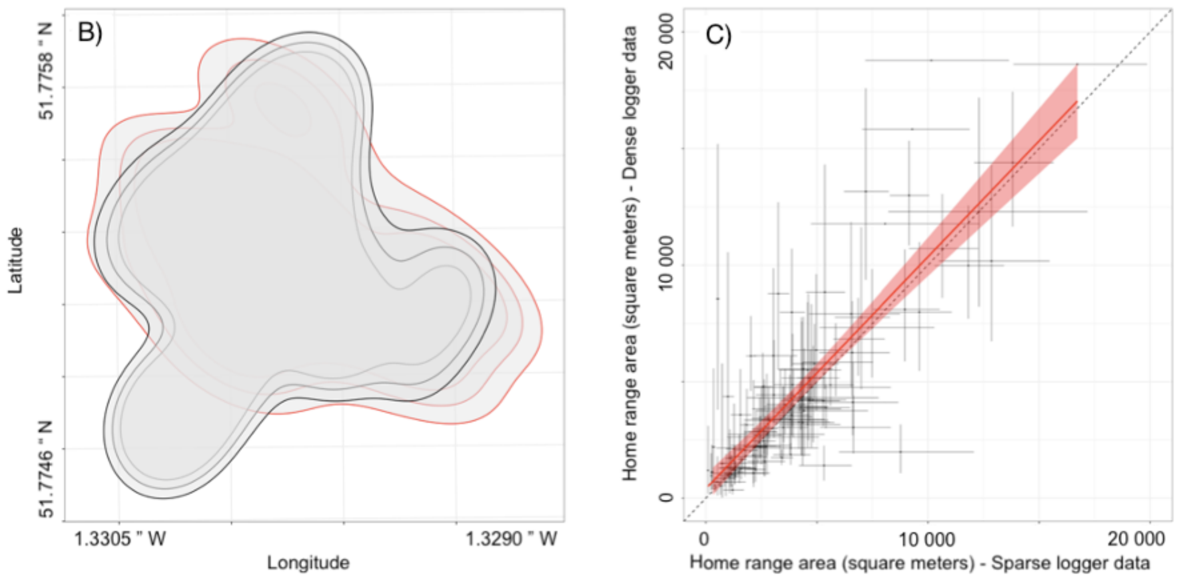


**Appendix S1 – Figure B.** Example of overlap between the home range of a single mouse constructed from sparser (red) vs denser (grey) logger data.

**Appendix S1 – Figure C.** Correlation between home range area estimates based on sparser (x-axis) and denser (y-axis) logger data, and the regression line (red line with confidence interval) from a linear model predicting one with the other. The area estimate combinations (points) have both horizontal and vertical confidence intervals (black lines).

#### Appendix S2. Details on gut microbiota analysis

For ease, the figure mentioned in this appendix is included in the end of this appendix and referred to as Figure A.

##### *Sample analysis order*

The microbiota samples were analysed in two batches, each including i) DNA extraction, ii) library preparation and iii) sequencing on Illumina MiSeq platform. Both batches included soil samples as well as faecal samples of two rodent species (Wood mouse, *Apodemus sylvaticus;* Yellow-necked mouse*, Apodemus flacvicollis*). The first batch had additionally faecal samples from bank voles (*Myodes glaerolus*). All samples were randomised across 96-well plates for DNA extraction and re-ordered for 96-well plates for library preparation to enable detection of extraction and PCR-related technical effects on the final data. Sequencing batches had a slight temporal difference: First batch contained only samples from the first half of the study (February-July), while the second batch contained samples from throughout the study (February-November).

##### *DNA extraction*

DNA was extracted from samples with a 96-well plate format Zymo Quick-DNATM Fecal/Soil Microbe kits (Zymo) according to manufacturer's instructions, using a Qiagen TissueLyser plate-format bead beater for DNA homogenization. Each DNA extraction plate contained one mock community (ZYMOBiomics Microbial Community Standard cat. No.D6300) to evaluate extraction, PCR and sequencing accuracy. After extraction, DNA concentrations in samples and controls were quantified with a Qubit 3.0 Fluorometer to verify extraction success. Control samples were additionally amplified for bacterial DNA with a 40-cycle PCR (see below) and product run on gel to make sure no bacterial DNA amplified from negative controls. No bands were seen on these gels.

##### *Library preparation and sequencing*

Extracted DNA samples were re-ordered for library preparation plates and amplified in a two-step PCR with primers 515F and 926R targeting an approximately 370 bp sequence in the V4-V5 region of the bacterial 16S rRNA gene (Walters et al., 2016). Each plate contained 95 samples and one negative PCR-control. First round PCR had 20µl reaction volume, consisting of: 10µl KAPA 2x Mastermix (KAPA Biosystems), 0.25µl each primer at 10µM, 4.5µl ultra pure water and 5µl extracted DNA. Cycling conditions were as follows: denaturation at 98°C for 2min, 10 cycles of 95°C for 20s, 65°C for 15s, 70°C for 45s, followed by a final extension at 72°C for 5min and a 4°C hold. To reduce PCR-induced variation, this first round PCR was done in duplicates after which the products pooled (by mixing 10µl +10µl of each product on a clean plate) and purified on AMPure magnetic bead purification plate (according to manufacturer's instructions). After purification, the second round PCR was prepared by mixing 9 µl of purified product with a 11 µl of master mix containing 10 µl KAPA Mastermix and 0.5+0.5 µl of forward and reverse Illumina Nextera barcoded indexing primers, added in unique combinations per each sample. The second PCR included 15 cycles with the same thermocycle conditions and was followed by the same AMPure purification step as the first PCR. Finally, amplified, indexed and purified 16S libraries were quantified for concentration (with Qubit), diluted to even concentration of 3-5 ng/ul and pooled for sequencing. Of each library preparation plate, the negative PCR control and 7 real samples (in diagonal order) were run on agrose gel in the end to ensure amplification success and lack of PCR-contamination. Extraction controls were not re-run on gels after amplification. Pooled libraries (x2) were sent off to the Centre for Genomic Research in Liverpool, where each pool was size-selected using a Pippin Prep, by excluding fragments outside the expected range. Libraries were then sequenced on two separate sequencing runs using 2x250bp paired-end sequencing on an Illumina MiSeq.

*Bioinformatics and preprocessing of microbiota sequence data*

Raw sequence data from each of the two sequencing batches was separately demultiplexed into their original samples and then processed through the DADA2 pipeline (version 1.14)(Callahan et al., 2016). Here, *cutadapt* (Martin, 2011) was first used to determine optimal trimming length to remove primer and adapter sequences. Based on this, 24 base pairs from the beginning of each read were trimmed away using the trimLeft argument in FilterAndTrim function in DADA2. As part of this function, and following visual inspection of sequence quality, low- quality tails were also trimmed, leaving 290 bp for forward and 230 bp for reverse reads. Next, sequences were dereplicated (identical sequences combined and their quality scores merged), amplicon sequence variants (ASVs) were inferred using the DADA2 algorithm and reverse paired-end reads were merged. Reads that could not be merged or were of abnormal length (<365 bp or > 371 bp) were removed from the dataset.

After this step, data from the two sequencing batches were combined and processed together, using the R package *phyloseq* (McMurdie & Holmes, 2013). First, chimeric sequences were removed from the combined data and taxonomy was assigned to ASVs against the Silva Database (Quast et al., 2013; version 138). Following this, we used iNext package in R (Hsieh et al., 2016) to inspect rarefaction and sample completeness curves to determine a minimum threshold for sample read depth. We also found evidence of contamination in some of our extraction plates: There were bacterial DNA in our control samples and in some plates, there was a spatial autocorrelation signal in microbiota similarity among samples extracted in the same plate, i.e. samples extracted closer to each other were more likely to share contaminant ASVs. Contamination was much stronger in extraction controls, even ones that went through a clean (uncontaminated) PCR, implying that our DNA extraction protocol was vulnerable to contamination. This is a common issue with plate-format extraction kits, such as the ZYMO 96-well plate format faecal/soil DNA extraction kit we used (Minich et al., 2019).

This kind of contamination likely happens through spillage between samples during the lysis step, and is prone to increase “technical noise” in the microbiota data. While this noise is random across biological groups in our data (as our samples were randomised across extraction plates, it can potentially diminish or mask some real biological differences between samples. We decided to mitigate this by

1) Subsetting data to contain only wood mouse faecal or soil samples and the remaining pre-processing steps were completed separately for these sample types.
2) Identifying potential contaminant ASVs with *decontam* algorithm (Davis et al., 2018) and filtering these out of the datasets.
3) Adding extraction plate and within-plate extraction distance as covariates in the models of microbiota similarity.

After this contamination mitigation step, we proceeded with processing the data by:

4) Removing singletons and doubletons, i.e. very rare taxa only present in abundance of <3 reads within sample,
5) Based on iNEXT completeness and rarefaction curves samples below read depth threshold (6000 reads for fecal samples, 800 for soil samples) were dropped out of the data as incomplete and unrepresentative.
6) Filtering away all ASVs assigned to Phylum “Cyanobacteria” or Family ”Mitochondria”, because these ASVs are likely not gut microbes but rather remnants of host DNA or plant chloroplasts from diet.
7) normalizing the read count data per sample as proportions.

The original number of unique sequence variants (ASVs) inferred by DADA2 algorithm was 6174 for wood mouse samples. After decontaminating the data, removing singletons and doubletons and all ASVs assigned to Cyanobacteria or Mitochondria, wood mouse samples had 1289 ASVs left.

Visualization of mock community composition revealed that our laboratory- and bioinformatics pipeline successfully captured the microbial diversity present in mock samples, though relative abundances varied somewhat from their theoretically true levels (Fig A below).

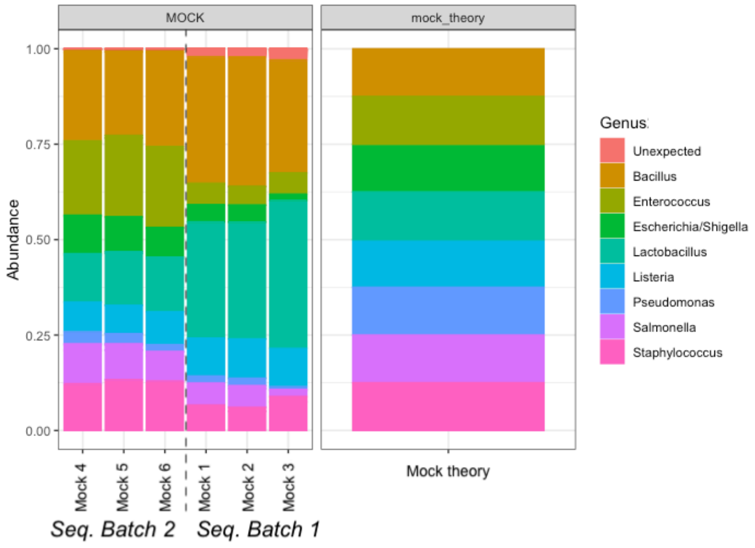


**Appendix S2 – Figure A. Microbial standard (mock) community profiles.**

Community composition in sequenced mock community samples compared to the expected composition (Mock theory).

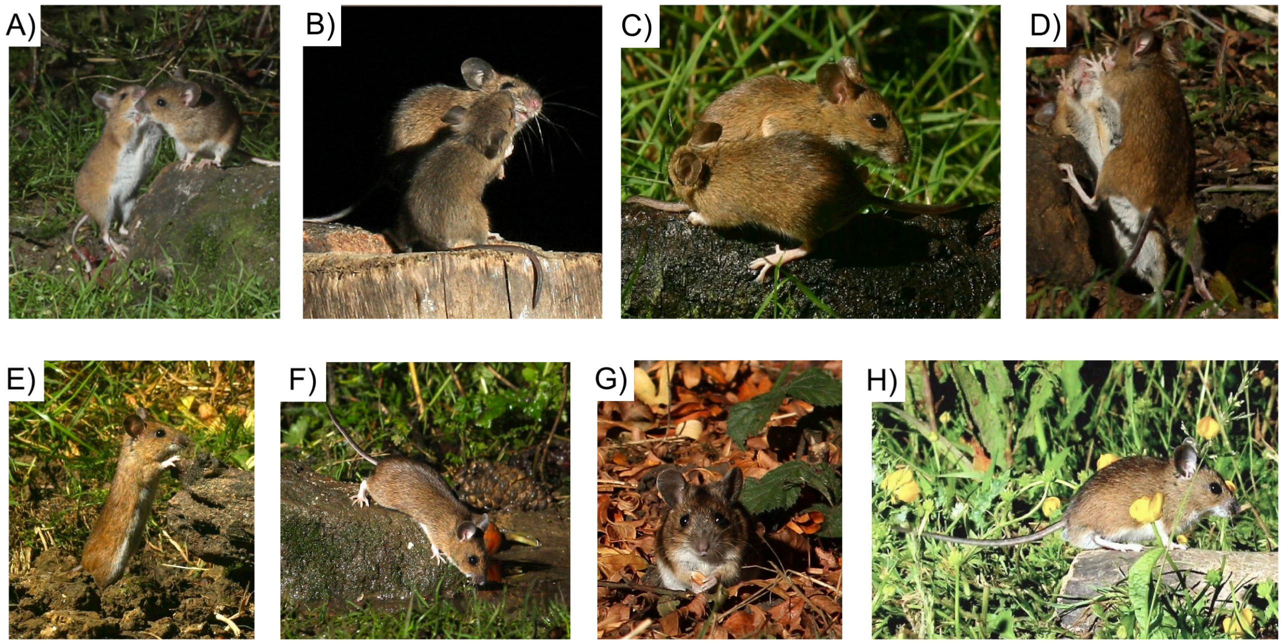


**Figure S1. Examples of social and environmental contact behaviours in wood mice.** Social contact behaviours that could transmit gut microbes include licking and grooming (A-B), anogenital inspections (C) and aggressive interactions (D). Individual variation in environmental contacts that could transmit gut microbes include ranging behaviour and exposure to different microhabitats and soils (E-H). These photos were taken with a motion sensor camera trap (with a light) during natural wood mouse activity hours (night) from a natural wood mouse habitat in Milton Keynes, UK. Photo credit Roy and Marie Battell.

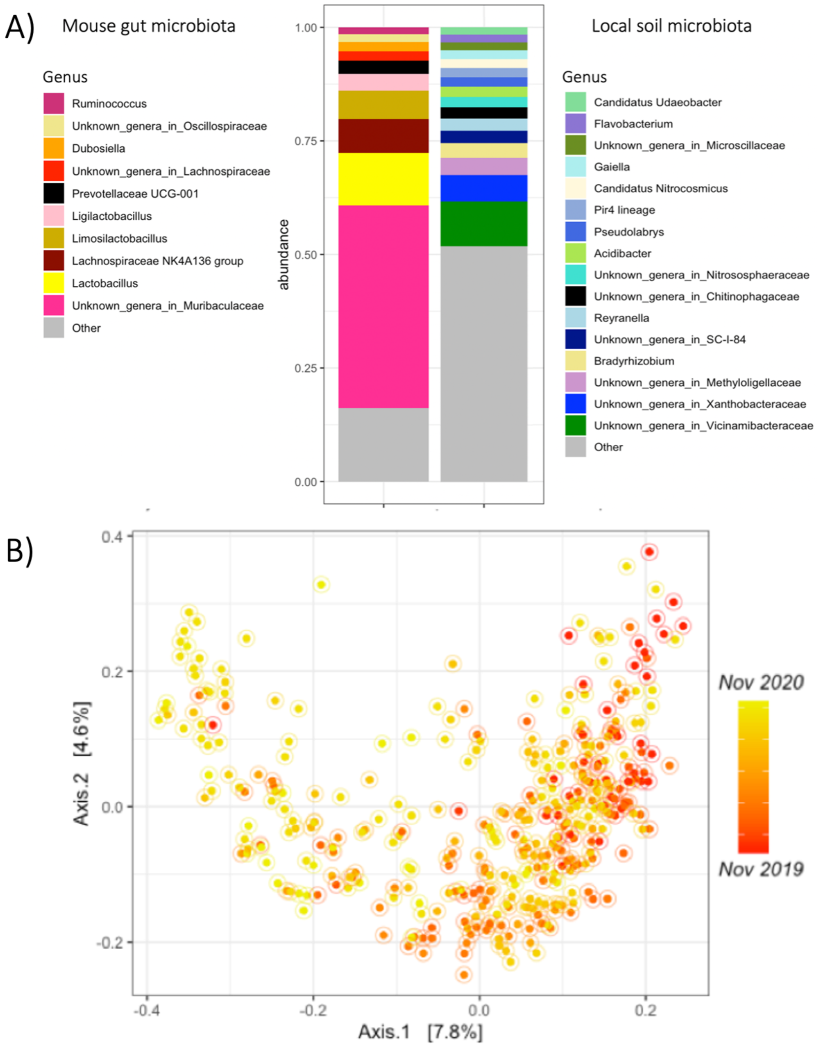


**Figure S2. Composition and variation in wood mouse gut microbiota and the local soil microbiota.** A) Taxonomic composition of the most abundant (>1%) genera in the wood mouse gut microbiota (left) and local soil microbiota (right). B) PCoA plot shows that the first axis of variation in wood mouse gut microbiota composition (Jaccard Index) reflects temporal changes in microbiota composition across the study period.

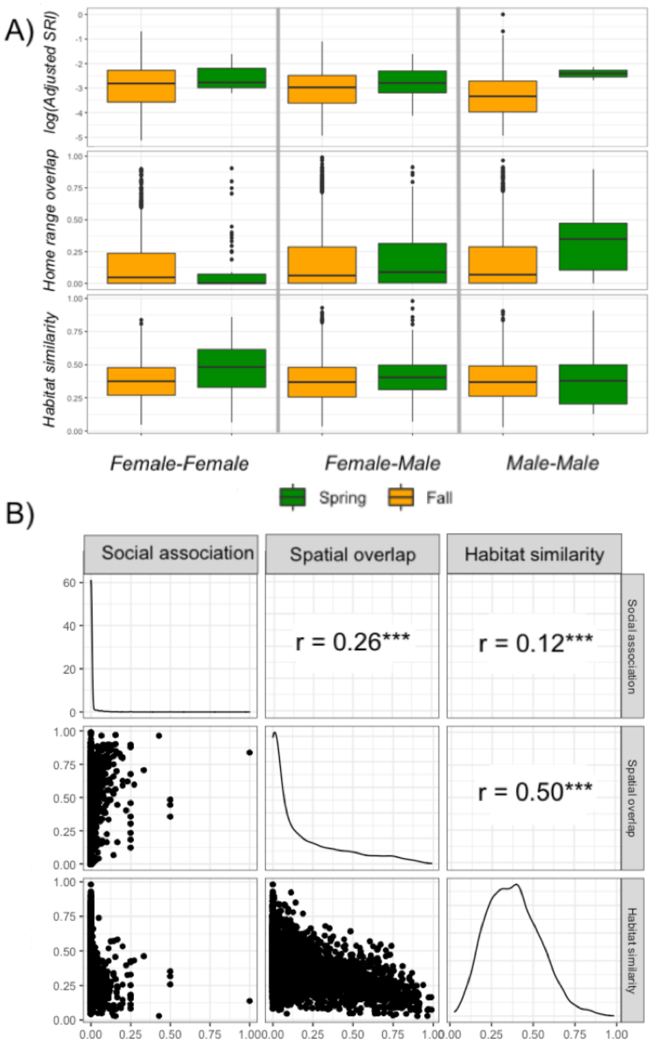


**Figure S3. Variation and covariation among social association, spatial overlap and habitat similarity.** A) Distribution of social association (upper panel), spatial overlap (middle panel) and habitat similarity (bottom panel) values across categories of sex-combinations (x-axis) and seasons (color). Heavily left-skewed values of social association values are reported on log-scale for ease of comparison. B) Correlation plot depicting associations among measures of social association (Adjusted SRI), spatial overlap (Bhattacaryya overlap between 75%? kernel utilization distributions) and habitat similarity (Bray-Curtis habitat similarity). Lower triangle illustrates covariation among the two intersecting variables, diagonal depicts the distribution of raw values in the intersecting variable and upper triangle reports the correlation among the intersecting variables, as measured by Mantel test statistic.

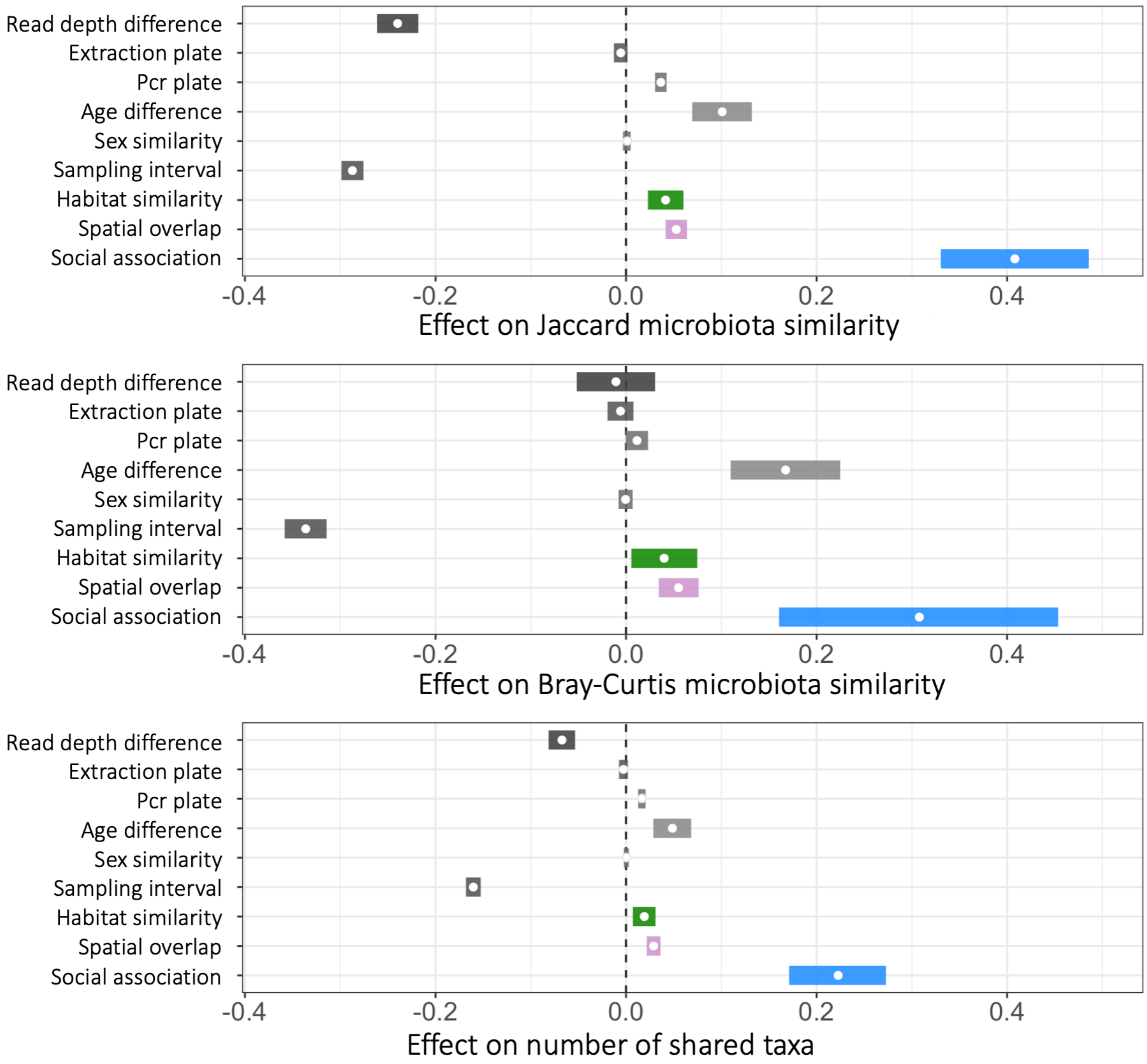


**Figure S4. Effects on alternative measures of microbiota similarity.**

Results from dyadic Bayesian regression models (brms) predicting Jaccard microbiota similarity (top panel), Abundance-weighted Bray-Curtis microbiota similarity (middle panel) and number of shared taxa (bottom panel). Posterior means (points) and their 95% credible intervals (coloured lines) are plotted from models (Supplementary Table S2B-C) with three alternative measures of pairwise microbiota similarity as the response. Where credible intervals do not overlap zero, a variable significantly predicts microbiota similarity while controlling for all other terms shown.

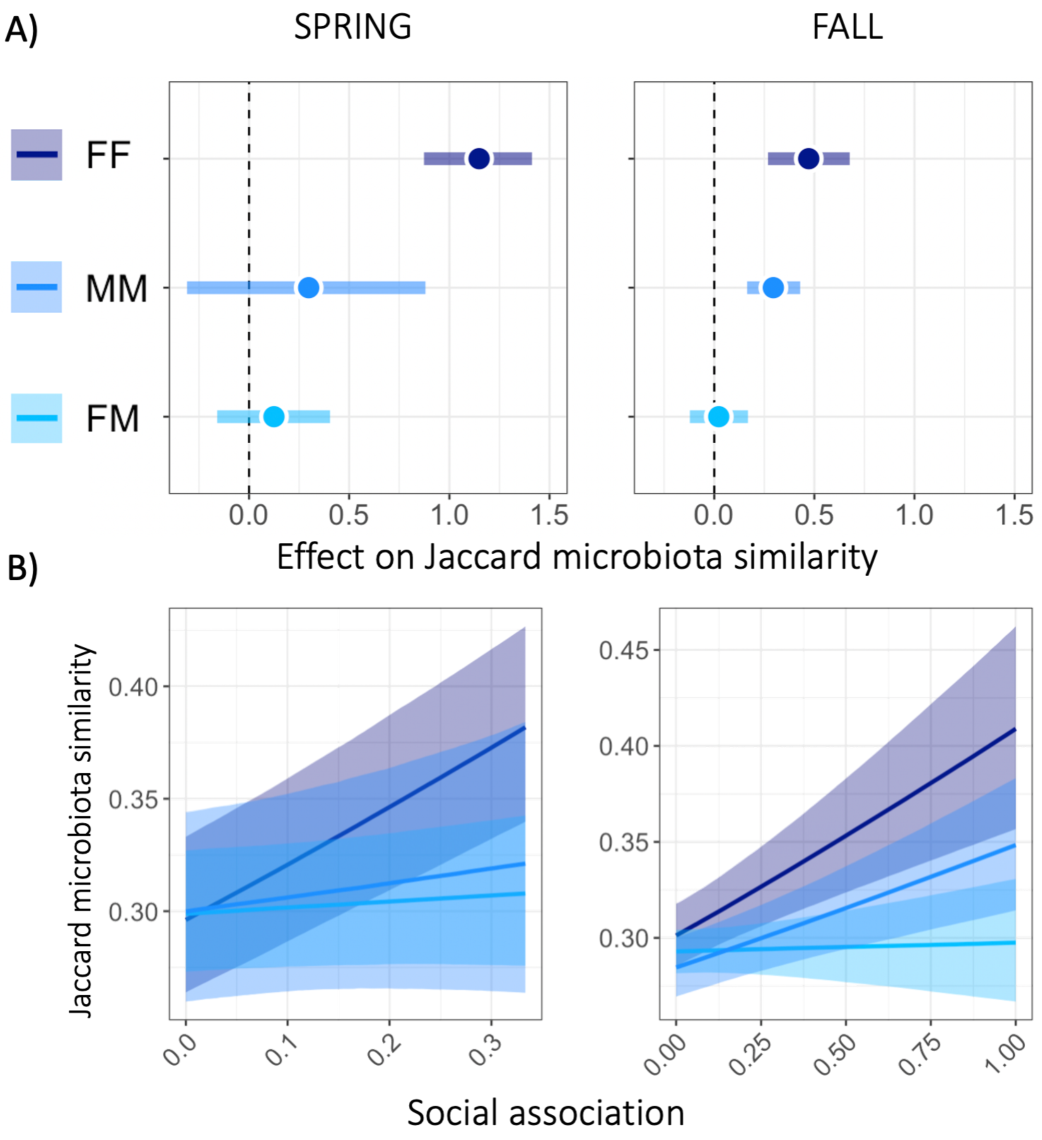


**Figure S5. Effects of social association on microbiota similarity across seasons and sexes.** Social effect on Jaccard microbiota similarity (x-axis) in pairs with different sex-combinations (colours: FF=female-female, MM=male-male, FM=female-male), in spring (left panel) vs. fall (right panel). Posterior means (points) and their 95% credible intervals (coloured lines) are plotted from Bayesian regression (brms) models (Supplementary Tables S4A-B). Where credible intervals do not overlap zero, a social association significantly predicts microbiota similarity. B) Slopes of social association effect on microbiota similarity across sex combinations (colours) and seasons (panels).

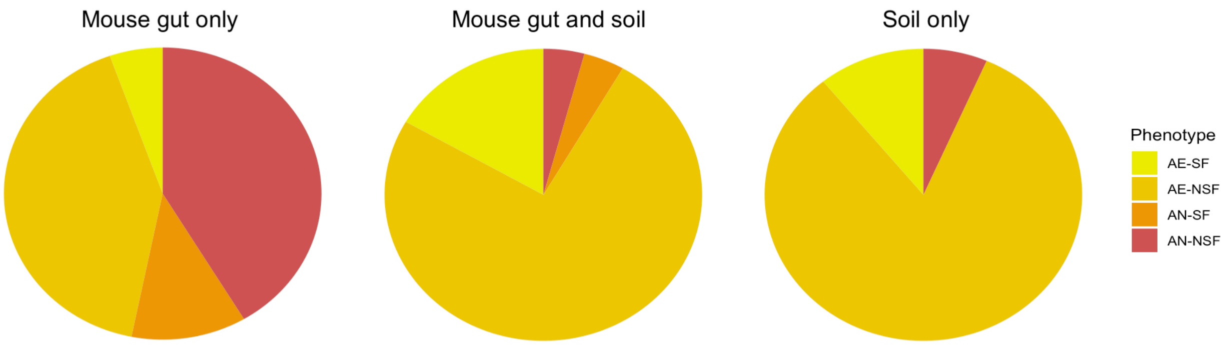


**Figure S6. Phenotype distribution among bacterial genera found in mouse gut, soil or both.** Proportions of Aerobe spore-formers (AE-SF, yellow), aerobe non-spore-formers (AE-NSF, gold), anaerobe spore-formers (AN-SF, orange) and anaerobe non-spore-formers (AN-NSF, red) among genera with available phenotype info found i) only in mouse gut (n=111, left chart), ii) both in mouse gut and soil (n=24, middle chart), and iii) only in prevalent soil-only microbiota (n=46, right panel). Prevalent soil-only microbiota here comprises of a subset of all soil-only taxa, specifically the genera found from at least 50% of soil samples and is meant to serve as a representation of taxa common in the soil.

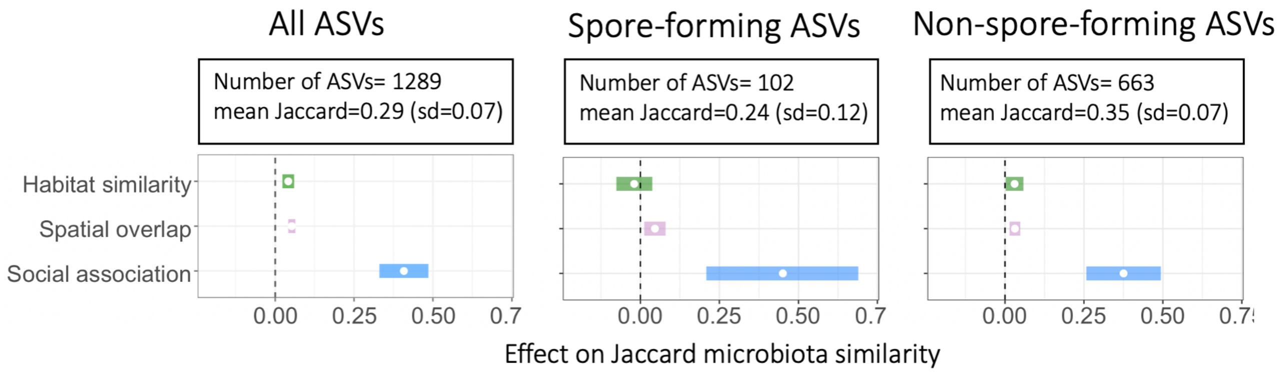


**Figure S7. Effects of main predictors on microbiota similarity in different phenotypic subsets of the microbiota – spore-formation**. Effects of social association (blue), spatial overlap (purple) and habitat similarity (green) on microbiota similarity among all ASVs (left), spore-forming ASVs (middle) and non-spore-forming ASVs (right). Posterior means (points) and their 95% credible intervals (coloured lines) are plotted from Bayesian regression (brms) models (Supplementary Table S*X*) with pairwise microbiota similarity among hosts (Jaccard Index) as the response. Where credible intervals do not overlap zero, a variable significantly predicts microbiota similarity while controlling for all other terms shown.

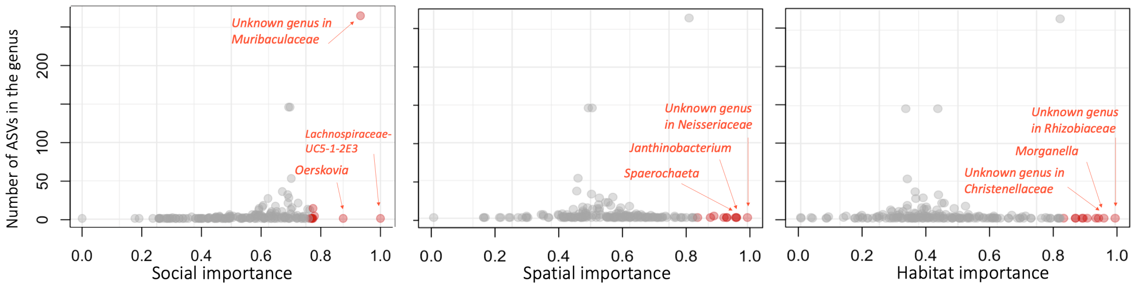


| Genus | Phylum | no. ASVs | Importance |
|---|---|---|---|
| Lachnospiraceae-UC5-1-2E3 | Firmicutes | 1 | 1.00 |
| Unknown genus in Muribaculaceae | Bacteroidota | 265 | 0.93 |
| Oerskovia | Actinobacteriota | 1 | 0.87 |
| Lachnospiraceae-A2 | Firmicutes | 4 | 0.78 |
| Ruminococcus | Firmicutes | 14 | 0.77 |
| Blastocatella | Acidobacteriota | 1 | 0.77 |
| Candidatus Soleaferrea | Firmicutes | 1 | 0.77 |
| Allobaculum | Firmicutes | 1 | 0.77 |
| Gordonibacter | Actinobacteriota | 1 | 0.77 |
| Colidextribacter | Firmicutes | 12 | 0.76 |
| Mannheimia | Proteobacteria | 1 | 0.76 |
| Mesorhizobium | Proteobacteria | 1 | 0.76 |
| Lactobacillus | Firmicutes | 13 | 0.76 |
| Novosphingobium | Proteobacteria | 1 | 0.75 |
| Unknown genus in Thermomicrobiales | Chloroflexi | 1 | 0.74 |
| Anaerostipes | Firmicutes | 4 | 0.74 |
| Prevotellaceae Ga6A1 group | Bacteroidota | 3 | 0.74 |
| Vitreoscilla | Proteobacteria | 1 | 0.74 |
| Curtobacterium | Actinobacteriota | 1 | 0.74 |
| Eubacterium brachy group | Firmicutes | 3 | 0.73 |

| Genus | Phylum | no. ASVs | Importance |
|---|---|---|---|
| Unknown genus in Neisseriaceae | Proteobacteria | 1 | 1.00 |
| Janthinobacterium | Proteobacteria | 1 | 0.97 |
| Sphaerochaeta | Spirochaetota | 1 | 0.97 |
| Microvirga | Proteobacteria | 1 | 0.96 |
| Curtobacterium | Actinobacteriota | 1 | 0.93 |
| Vitreoscilla | Proteobacteria | 1 | 0.93 |
| Elusimicrobium | Elusimicrobiota | 1 | 0.92 |
| Rikenellaceae RC9 gut group | Bacteroidota | 3 | 0.89 |
| Unknown genus in Rhizobiaceae | Proteobacteria | 1 | 0.88 |
| Unknown genus in Coriobacteriales | Actinobacteriota | 1 | 0.84 |
| Variovorax | Proteobacteria | 1 | 0.83 |
| Pantoea | Proteobacteria | 1 | 0.83 |
| Peptococcus | Firmicutes | 1 | 0.83 |
| Unknown genus in Erysipelatoclostridiaceae | Firmicutes | 1 | 0.82 |
| Unknown genus in Muribaculaceae | Bacteroidota | 265 | 0.81 |
| Oerskovia | Actinobacteriota | 1 | 0.81 |
| Moraxella | Proteobacteria | 1 | 0.79 |
| Ferruginibacter | Bacteroidota | 1 | 0.79 |
| Puia | Bacteroidota | 1 | 0.79 |
| Polaromonas | Proteobacteria | 1 | 0.77 |

| Genus | Phylum | no. ASVs | Importance |
|---|---|---|---|
| Unknown genus in Rhizobiaceae | Proteobacteria | 1 | 1.00 |
| Morganella | Proteobacteria | 1 | 0.96 |
| Unknown genus in Christensenellaceae | Firmicutes | 1 | 0.95 |
| Candidatus Saccharimonas | Patescibacteria | 1 | 0.94 |
| Pantoea | Proteobacteria | 1 | 0.91 |
| Bilophila | Desulfobacterota | 1 | 0.90 |
| Allobaculum | Firmicutes | 1 | 0.90 |
| Polaromonas | Proteobacteria | 1 | 0.87 |
| Mannheimia | Proteobacteria | 1 | 0.87 |
| Acidibacter | Proteobacteria | 1 | 0.84 |
| Unknown genus in Muribaculaceae | Bacteroidota | 265 | 0.82 |
| Nakamurella | Actinobacteriota | 1 | 0.82 |
| Nocardioides | Actinobacteriota | 1 | 0.82 |
| Oscillospiraceae-UCG-005 | Firmicutes | 1 | 0.82 |
| Massilia | Proteobacteria | 1 | 0.82 |
| Paraprevotella | Bacteroidota | 1 | 0.82 |
| Amaricoccus | Proteobacteria | 1 | 0.79 |
| Ochroba trum | Proteobacteria | 1 | 0.79 |
| Candidatus Soleaferrea | Firmicutes | 1 | 0.79 |
| Actinoplanes | Actinobacteriota | 2 | 0.78 |

**Figure S8. Overview of the most important genera for social, spatial and habitat signal in microbiota similarity.** Top panel: Distribution of social (left), spatial (middle) and habitat (right) importance scores (x-axis) against the number of ASVs in each genus (y-axis). Top 10 most important genera are marked red. Bottom table: Overview of top 20 most important genera in each importance type. Here too, top 10 most important genera in each category are marked red.

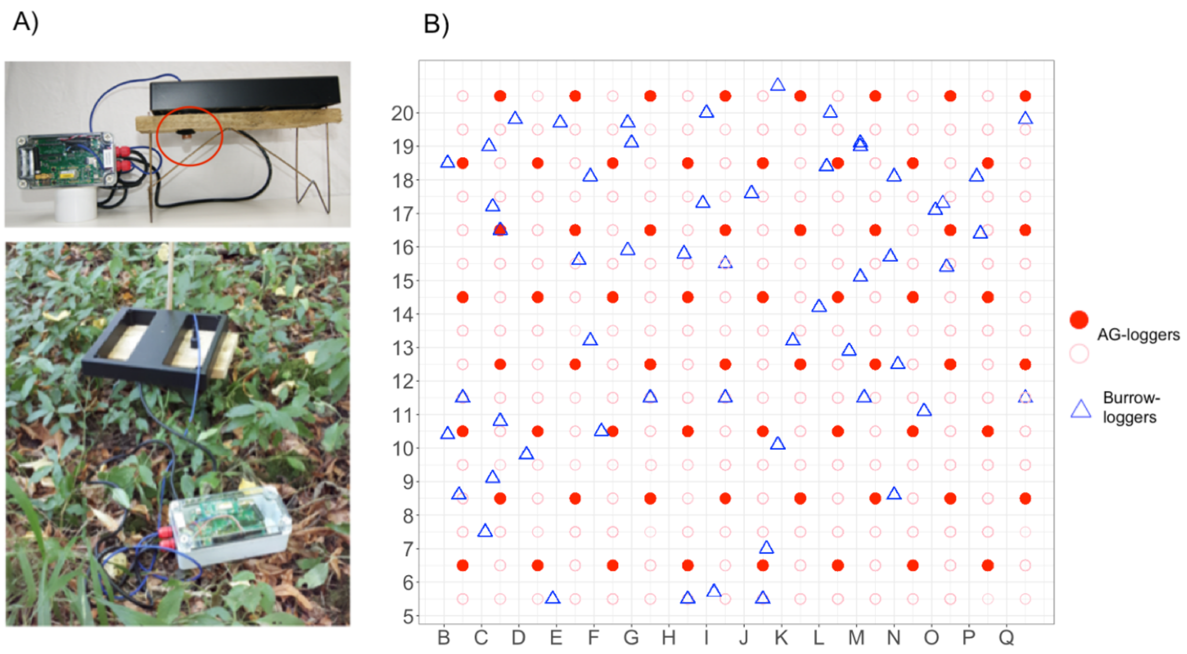


### Figure S9. The RFID logger device and tracking protocol

A) The logger was un-baited and consisted of a motion sensor (red circle, top figure), that woke up the detection coil (black square) whenever a warm object, such as a rodent came nearby. The coil, sitting on a wooden "table" to allow rodents to pass under as well as around it, then created a ~1 m2 oscillating detection field that would interfere – and subsequently log- any 124 kHz RFID tag within this field. Exploiting this, wild rodents were tagged with a subcutaneous passive integrated transponder (PIT-) tags containing a 10-digit HEX-code identification number readable in this exact oscillation frequency. When a rodent tag was read by the coil, this identification number was saved on an SD-card (in the grey box) and additionally transmitted into and interactive internet database through an antenna (in the grey box) connected to the broadband network covering Wytham Woods (Wytham Data Net). B) 60 loggers (red dots) were positioned in an even chequerboard design, where loggers would be in the middle of every fourth 10 x 10 m grid cell across the 2.56 study grid. Additionally, loggers were fortnightly rotated throughout the whole study (February-November 2019) by moving each logger one cell (=10 m) north (increasing numbers in y axis). This way each grid cell had a logger (pink circles) for two weeks every two months.

In addition to these 60 loggers, for a small subset of analyses, we used data from an extra set of 60 Burrow-loggers (blue triangles), which were positioned approximately evenly across the grid, on top of a set of known active mouse burrows across the grid from July until November, and not rotated. Burrow-logger data was used to complement AG-logger data in home-range estimation, but not in any other logger-derived measures.

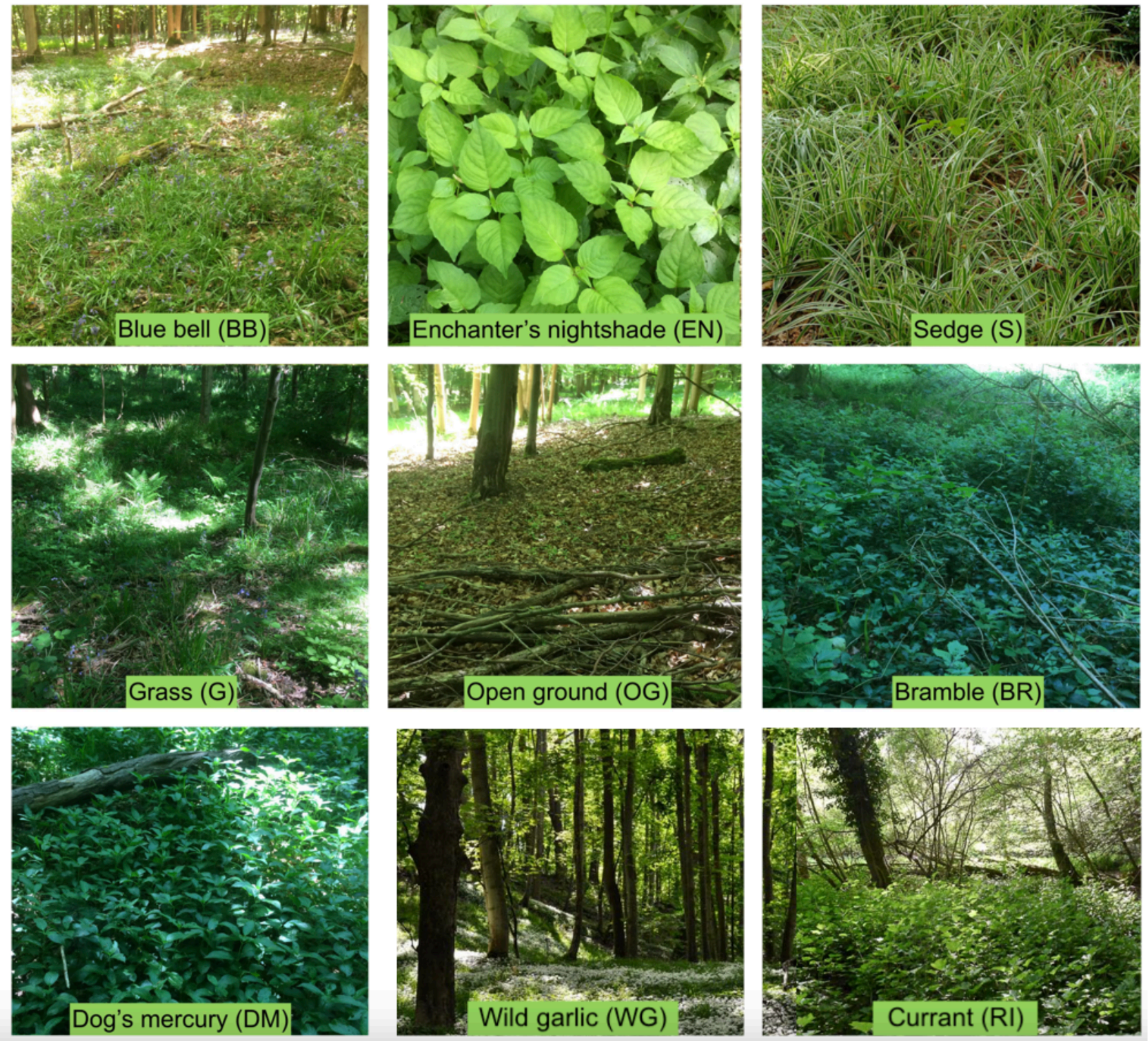


**Figure S10**: **Main ground cover habitat types in Holly Hill study site.**

Photos are taken from Wytham Woods by authors except the "currant"-landscape which is from a similar woodland in Sussex, England (Photo credit Paul Kirtley).

#### Table S1. Model results: Effects of Social association, spatial overlap and habitat similarity on microbiota similarity.

Results of *brms* models testing the effect of social association, spatial overlap, habitat similarity and covariates on different measures of microbiota similarity: A) Jaccard Index (betaregression), B) Bray-Curtis similarity (betaregression), C) number of shared taxa (poisson regression). Significant terms (where 95% credible intervals do not include zero) are shown in bold. Est.Error indicates the standard deviation of the posterior distribution.

| **Table S1A. Predicting Jaccard microbiota similarity** | | | | |
|---|---|---|---|---|
| | Estimate | Est.Error | l-95% CI | u-95% CI |
| Intercept | -0.93 | 0.03 | -0.99 | -0.88 |
| **Sample read depth difference** | -0.24 | 0.01 | -0.26 | -0.22 |
| Extraction distance | -0.01 | 0.00 | -0.01 | 0.00 |
| **PCR-plate similarity** | 0.04 | 0.00 | 0.03 | 0.04 |
| **Age-similarity** | 0.10 | 0.02 | 0.07 | 0.13 |
| Sex similarity | 0.00 | 0.00 | -0.00 | 0.00 |
| **Sampling interval** | -0.29 | 0.01 | -0.30 | -0.28 |
| **Habitat similarity** | 0.04 | 0.01 | 0.02 | 0.06 |
| **Spatial overlap** | 0.05 | 0.01 | 0.04 | 0.06 |
| **Social association** | 0.41 | 0.04 | 0.33 | 0.49 |
| **Table S1B. Predicting Bray-Curtis microbiota similarity** | | | | |
| | Estimate | Est.Error | l-95% CI | u-95% CI |
| Intercept | -1.09 | 0.05 | -1.19 | -1.00 |
| **Sample read depth difference** | -0.01 | 0.02 | -0.05 | 0.03 |
| Extraction distance | -0.01 | 0.01 | -0.02 | 0.01 |
| **PCR-plate similarity** | 0.01 | 0.01 | -0.00 | 0.02 |
| **Age-similarity** | 0.17 | 0.03 | 0.11 | 0.22 |
| Sex similarity | -0.00 | 0.00 | -0.01 | 0.01 |
| **Sampling interval** | -0.34 | 0.01 | -0.36 | -0.31 |
| **Habitat similarity** | 0.04 | 0.02 | 0.01 | 0.07 |
| **Spatial overlap** | 0.05 | 0.01 | 0.03 | 0.08 |
| **Social association** | 0.31 | 0.08 | 0.16 | 0.45 |
| **Table S1C. Predicting number of shared taxa** | | | | |
| | Estimate | Est.Error | l-95% CI | u-95% CI |
| Intercept | 4.36 | 0.03 | 4.30 | 4.42 |
| **Sample read depth difference** | -0.07 | 0.01 | -0.08 | -0.05 |
| Extraction distance | -0.00 | 0.00 | -0.01 | 0.00 |
| **PCR-plate similarity** | 0.02 | 0.00 | 0.01 | 0.02 |

| | | | | |
|---|---|---|---|---|
| **Age-similarity** | 0.05 | 0.01 | 0.03 | 0.07 |
| Sex similarity | 0.00 | 0.00 | -0.00 | 0.00 |
| **Sampling interval** | -0.16 | 0.00 | -0.17 | -0.15 |
| **Habitat similarity** | 0.03 | 0.01 | 0.01 | 0.03 |
| **Spatial overlap** | 0.02 | 0.00 | 0.02 | 0.04 |
| **Social association** | 0.22 | 0.03 | 0.17 | 0.27 |

#### Table S2. Model results: Effects of social association, spatial overlap and habitat similarity on microbiota similarity among pairs with different sex categories.

Results of *brms* models testing whether effects of social association, spatial overlap and habitat similarity on microbiota similarity (Jaccard Index) varied according to the sex combination of mouse pairs, while accounting for covariates. Significant terms are those where 95% credible intervals do not include zero. Est.Error indicates the standard deviation of the posterior distribution. Interaction effects are reported as slopes relative to the base category of female-female as per model output.

**Table S2. Effects on microbiota similarity across sex combinations**

| | Estimate | Est.Error | l-95% CI | u-95% CI |
|---|---|---|---|---|
| Intercept | -0.90 | 0.04 | -0.97 | -0.82 |
| Sample read depth difference | -0.24 | 0.01 | -0.26 | -0.22 |
| Extraction distance | -0.01 | 0.00 | -0.01 | 0.00 |
| PCR-plate similarity | 0.04 | 0.00 | 0.03 | 0.04 |
| Age similarity | 0.10 | 0.02 | 0.07 | 0.13 |
| Sampling interval | -0.29 | 0.01 | -0.30 | -0.28 |
| Habitat similarity | 0.02 | 0.02 | -0.01 | 0.05 |
| Sex category (female-male) | -0.03 | 0.03 | -0.08 | 0.03 |
| Sex category (male-male) | -0.10 | 0.05 | -0.20 | 0.00 |
| Spatial overlap | 0.10 | 0.01 | 0.07 | 0.12 |
| Social association | 0.74 | 0.08 | 0.59 | 0.90 |
| Habitat similarity: Sex category (female-male) | 0.00 | 0.02 | -0.03 | 0.04 |
| Habitat similarity: Sex category (male-male) | 0.07 | 0.02 | 0.02 | 0.11 |
| Spatial overlap: Sex category (female-male) | -0.05 | 0.01 | -0.08 | -0.02 |
| Spatial overlap: Sex category (male-male) | -0.07 | 0.02 | -0.10 | -0.04 |
| Social association: Sex category (female-male) | -0.56 | 0.10 | -0.76 | -0.37 |
| Social association: Sex category (male-male) | -0.34 | 0.10 | -0.54 | -0.14 |

#### Table S3. Model results: Effects of spring and fall social association on microbiota similarity among pairs with different sex categories.

Results of *brms* models testing the effects of spring and fall social association across sex combinations on microbiota similarity (Jaccard Index), while accounting for covariates. Age is not included as a covariate in the spring model, as there was not enough age variation in the spring (only adults). Significant terms (where 95% credible intervals do not include zero) are shown in bold. Est.Error indicates the standard deviation of the posterior distribution. For ease of interpretation, interaction effects between sex combination and social association are *NOT* reported as slopes relative to each other but as independent effects within each group (e.g., "Male-male Spring social association" means the independent effect of social association on microbiota among males).

| **Table S3A. Spring** | | | | |
|---|---|---|---|---|
| | Estimate | Est.Error | l-95% CI | u-95% CI |
| Intercept | -0.81 | 0.09 | -0.97 | -0.64 |
| **Sample read depth difference** | -0.21 | 0.04 | -0.30 | -0.13 |
| Extraction distance | 0.00 | 0.01 | -0.02 | 0.02 |
| **PCR-plate similarity** | 0.02 | 0.01 | 0.00 | 0.04 |
| **Sampling interval** | -0.52 | 0.03 | -0.57 | -0.46 |
| **Habitat similarity** | 0.14 | 0.03 | 0.07 | 0.21 |
| Spatial overlap | -0.02 | 0.02 | -0.07 | 0.02 |
| Sex category (female-male) | 0.01 | 0.07 | -0.12 | 0.14 |
| Sex category (male-male) | 0.02 | 0.13 | -0.25 | 0.28 |
| **Female-female Spring social association** | 1.15 | 0.14 | 0.88 | 1.42 |
| Female-male Spring social association | 0.12 | 0.19 | -0.16 | 0.40 |
| Male-male Spring social association | 0.30 | 0.32 | -0.31 | 0.88 |
| **Table S3B. Fall** | | | | |
| | Estimate | Est.Error | l-95% CI | u-95% CI |
| Intercept | -0.91 | 0.04 | -1.00 | -0.83 |
| **Sample read depth difference** | -0.24 | 0.01 | -0.27 | -0.21 |
| Extraction distance | -0.01 | 0.00 | -0.01 | 0.00 |
| **PCR-plate similarity** | 0.03 | 0.00 | 0.03 | 0.04 |
| **Age similarity** | 0.11 | 0.02 | 0.08 | 0.14 |
| **Sampling interval** | -0.21 | 0.02 | -0.24 | -0.19 |
| **Habitat similarity** | 0.03 | 0.01 | 0.01 | 0.06 |
| **Spatial overlap** | 0.10 | 0.01 | 0.08 | 0.11 |
| Sex category (female-male) | -0.04 | 0.03 | -0.09 | 0.01 |
| Sex category (male-male) | -0.08 | 0.05 | -0.19 | 0.03 |
| **Female-female Fall social association** | 0.47 | 0.10 | 0.27 | 0.68 |
| Female-male Fall social association | 0.02 | 0.12 | -0.12 | 0.17 |
| **Male-male Fall social association** | 0.30 | 0.12 | 0.16 | 0.43 |

#### Table S4. Aerotolerance and spore-forming ability of bacterial genera

Table is attached as a separate csv file. We used Bergey's Manual of Systematics of Archaea and Bacteria alongside other individual scientific publications to find information about the aerotolerance and spore-formation abilities of bacterial genera present in our data. References mentioned in the table are included in the Supplementary References in the end of this document. Specifically, we searched information for the 188 genera present in the mouse gut or mouse gut and soil and additionally 111 genera present only in the soil. For these soil-only taxa, we only considered those which were present in at least half of the soil samples. For genera with mixed or unknown aerotolerance or sporulation, we labelled "unknown". For unknown genera within a known family, we were conservative in that we generally considered them as having unknown aerotolerance and spore-formation. However, when other members of the family were consistently of a given phenotype, we gave a genus the phenotype of its family. For example, since all described strains in the family Muribaculaceae are anaerobic and non-spore-forming, we considered "unknown genus in Muribaculaceae" to also be anaerobic and non-spore-forming. For some genera with missing information on these phenotypic traits, we also labelled them after another name for the same genus present in the literature (e.g., "UBA1819" as "Faecalibacterium") or according to the genus that they had been previously classified as belonging to (e.g., *Lysinibacillus* as part of *Bacillus*). All genera assigned an aerotolerance or spore-formation label in any of these indirect manners, have been marked in the columns "Aerotolerance_indirectly_inferred" and "Sporeformation_indirectly_inferred", and the taxon after which they were labelled is specified in the "References" column. Spore-formation column labels taxa as spore-forming (SF) or non-spore-forming (NSF). Aerotolerance column labels taxa as anaerobic (=obligate anaerobes) or aerotolerant (all facultative anaerobes, microaerobes, or aerobes). As in the analyses, as in Suzuki et al., 2019, all taxa with any aerotolerance (all except obligate anaerobes) where considered aerotolerant.

#### Table S5. Model results: Effects on microbiota similarity in different phenotypic subsets of microbiota.

Results of brms models testing the effect of social association, spatial overlap, habitat similarity and covariates on gut microbiota similarity (Jaccard Index) of a) aerotolerant taxa or b) anaerobic taxa, c) spore-forming taxa and d) non-spore-forming taxa. Significant terms (where 95% credible intervals do not include zero) are shown in bold. Est.Error indicates the standard deviation of the posterior distribution. Note that posterior distributions are not directly comparable between models due to inherently different uncertainty, intercepts and slopes across phenotypic data subsets.

| **Table S5 Aerotolerant microbiota** | | | | |
|---|---|---|---|---|
| | Estimate | Est.Error | l-95% CI | u-95% CI |
| Intercept | -1.08 | 0.10 | -1.27 | -0.88 |
| **Sample read depth difference** | -0.42 | 0.04 | -0.50 | -0.34 |
| Extraction distance | -0.05 | 0.01 | -0.08 | -0.03 |
| **PCR-plate similarity** | 0.26 | 0.01 | 0.24 | 0.28 |
| Age-similarity | 0.10 | 0.06 | -0.03 | 0.22 |
| **Sex similarity** | -0.01 | 0.01 | -0.02 | 0.01 |
| **Sampling interval** | -0.76 | 0.02 | -0.81 | -0.72 |
| Habitat similarity | -0.03 | 0.04 | -0.10 | 0.04 |
| **Spatial overlap** | 0.07 | 0.02 | 0.03 | 0.11 |
| Social association | 0.17 | 0.15 | -0.12 | 0.46 |
| **Table S5B. Anaerobic microbiota** | | | | |
| | Estimate | Est.Error | l-95% CI | u-95% CI |
| Intercept | -1.17 | 0.04 | -1.25 | -1.09 |
| **Sample read depth difference** | -0.29 | 0.02 | -0.33 | -0.25 |
| Extraction distance | 0.00 | 0.01 | -0.01 | 0.02 |
| **PCR-plate similarity** | 0.02 | 0.01 | 0.01 | 0.03 |
| **Age-similarity** | 0.09 | 0.03 | 0.04 | 0.14 |
| Sex similarity | 0.00 | 0.00 | 0.00 | 0.01 |
| **Sampling interval** | -0.29 | 0.01 | -0.31 | -0.27 |
| **Habitat similarity** | 0.04 | 0.02 | 0.00 | 0.07 |
| **Spatial overlap** | 0.03 | 0.01 | 0.01 | 0.05 |
| **Social association** | 0.42 | 0.07 | 0.28 | 0.56 |
| **Table S5C Spore-forming microbiota** | | | | |
| | Estimate | Est.Error | l-95% CI | u-95% CI |
| Intercept | -1.24 | 0.08 | -1.40 | -1.08 |
| **Sample read depth difference** | -0.05 | 0.03 | -0.11 | 0.02 |
| Extraction distance | 0.00 | 0.01 | -0.03 | 0.02 |
| **PCR-plate similarity** | 0.00 | 0.01 | -0.01 | 0.02 |

| | | | | |
|---|---|---|---|---|
| Age-similarity | 0.05 | 0.05 | -0.05 | 0.14 |
| **Sex similarity** | 0.02 | 0.01 | 0.00 | 0.03 |
| **Sampling interval** | -0.22 | 0.02 | -0.25 | -0.18 |
| Habitat similarity | -0.02 | 0.03 | -0.08 | 0.04 |
| **Spatial overlap** | 0.05 | 0.02 | 0.01 | 0.08 |
| Social association | 0.45 | 0.12 | 0.21 | 0.69 |
| **Table S5D. Non-spore-forming microbiota** | | | | |
| | Estimate | Est.Error | l-95% CI | u-95% CI |
| Intercept | -1.01 | 0.04 | -1.08 | -0.94 |
| **Sample read depth difference** | -0.34 | 0.02 | -0.37 | -0.30 |
| Extraction distance | -0.01 | 0.01 | -0.02 | 0.00 |
| **PCR-plate similarity** | 0.06 | 0.00 | 0.05 | 0.07 |
| **Age-similarity** | 0.10 | 0.02 | 0.06 | 0.15 |
| Sex similarity | 0.00 | 0.00 | -0.01 | 0.01 |
| **Sampling interval** | -0.38 | 0.01 | -0.40 | -0.36 |
| **Habitat similarity** | 0.03 | 0.01 | 0.00 | 0.06 |
| **Spatial overlap** | 0.03 | 0.01 | 0.01 | 0.05 |
| **Social association** | 0.38 | 0.06 | 0.26 | 0.49 |

#### Table S6. Model results: Effects of Phenotypes on importance scores across bacterial genera.

Results of three brms models testing the effect of binary Aerotolerance and Sporulation (0/1) on importance scores of social (model S6A), spatial (model S6B) and habitat (model S6C) signal across bacterial genera, while controlling for their phylogeny. Significant terms (where 95% credible intervals do not overlap zero) are shown in bold. Est.Error indicates the standard deviation of the posterior distribution. Model formulas were:

Social Importance~ Aerotolerance*Sporulation + (1 | gr(genus, cov = Phylogenetic tree).

Spatial Importance~ Aerotolerance*Sporulation + (1 | gr(genus, cov = Phylogenetic tree).

Habitat Importance~ Aerotolerance*Sporulation + (1 | gr(genus, cov = Phylogenetic tree).

Here, interaction effects are reported as slopes relative to each other as per model output.

| **Table S6A. Social Importance** | | | | |
|---|---|---|---|---|
| | Estimate | Est.Error | l-95% CI | u-95% CI |
| Intercept | 0.58 | 0.06 | 0.43 | 0.67 |
| **Aerotolerance** | **-0.07** | **0.04** | **-0.14** | **-0.00** |
| Sporulation | -0.05 | 0.06 | -0.16 | 0.06 |
| Aerotolerance:Sporulation | 0.08 | 0.08 | -0.08 | 0.24 |
| **Table S6B. Spatial Importance** | | | | |
| | Estimate | Est.Error | l-95% CI | u-95% CI |
| Intercept | 0.48 | 0.10 | 0.24 | 0.61 |
| Aerotolerance | -0.02 | 0.04 | -0.10 | 0.06 |
| Sporulation | 0.02 | 0.06 | -0.10 | 0.13 |
| Aerotolerance:Sporulation | 0.13 | 0.08 | -0.03 | 0.28 |
| **Table S6C. Habitat Importance** | | | | |
| | Estimate | Est.Error | l-95% CI | u-95% CI |
| Intercept | 0.47 | 0.06 | 0.33 | 0.58 |
| Aerotolerance | -0.02 | 0.05 | -0.12 | 0.07 |
| **Sporulation** | **-0.17** | **0.07** | **-0.31** | **-0.02** |
| **Aerotolerance:Sporulation** | **0.21** | **0.10** | **0.00** | **0.41** |

**Table S7. Model results: Effects of microbial combination phenotypes on importance scores across bacterial genera.** Results of three brms models testing the effect of bacterial phenotype (4-level factor: aerotolerant spore-former AE-SF, aerotolerant non-spore-former AE-NSF, anaerobic spore-former AN-SF, anaerobic non-spore-former AN-NSF) on importance scores of social (model S7A), spatial (model S7B) and habitat (model S7C) signal across bacterial genera, while controlling for their phylogeny. Significant terms are those where 95% credible intervals do not include zero. Est.Error indicates the standard deviation of the posterior distribution. Model formulas were:

Social Importance ~ phenotype+ (1 | gr(genus, cov = Phylogenetic tree)

Spatial Importance ~ phenotype+ (1 | gr(genus, cov = Phylogenetic tree)

Habitat Importance ~ phenotype+ (1 | gr(genus, cov = Phylogenetic tree)

| **Table S7A: Social importance** | | | | |
|---|---|---|---|---|
| | Estimate | Est.Error | l-95% CI | u-95% CI |
| Intercept | 0.58 | 0.06 | 0.43 | 0.67 |
| Phenotype (AN-SF) | -0.05 | 0.06 | -0.16 | 0.07 |
| Phenotype (AE-NSF) | -0.07 | 0.04 | -0.15 | -0.00 |
| Phenotype (AE-SF) | -0.04 | 0.06 | -0.16 | 0.07 |
| **Table S7B: Spatial importance** | | | | |
| | Estimate | Est.Error | l-95% CI | u-95% CI |
| Intercept | 0.61 | 0.10 | 0.37 | 0.76 |
| Phenotype (AN-NSF) | -0.12 | 0.06 | -0.24 | -0.00 |
| Phenotype (AN-SF) | -0.10 | 0.08 | -0.25 | 0.04 |
| Phenotype (AE-NSF) | -0.14 | 0.06 | -0.25 | -0.03 |
| **Table S7C: Habitat importance** | | | | |
| | Estimate | Est.Error | l-95% CI | u-95% CI |
| Intercept | 0.48 | 0.06 | 0.36 | 0.59 |
| Phenotype (AN-SF) | -0.22 | 0.08 | -0.38 | -0.06 |
| Phenotype (AE-NSF) | -0.02 | 0.05 | -0.11 | 0.07 |
| Phenotype (AE-SF) | 0.05 | 0.08 | -0.11 | 0.21 |

**Table S8. Model results: Results of post hoc models testing whether specific phenotype categories differ significantly from other genera in their importance.** Results of three brms models, testing whether A) Anaerobe non-spore formers (AN-NSF) have significantly higher social importance than other genera, B) Aerobe-spore formers (AE-SF) have significantly higher spatial importance than other genera and C) Anaerobe spore formers (AN-SF) have significantly lower habitat importance than other genera. Predictor term was a binary variable describing whether a genus was of a given phenotype (1) or not (0). Significant terms (where 95% credible intervals do not include zero) are shown in bold. Est.Error indicates the standard deviation of the posterior distribution. Model formulas were:

Social Importance ~ AN-NSF + (1 | gr(genus, cov = Phylogenetic tree)

Spatial Importance ~ AE-SF + (1 | gr(genus, cov = Phylogenetic tree)

Habitat Importance ~ AN-SF + (1 | gr(genus, cov = Phylogenetic tree)

| **Table S8A: Social importance** | | | | |
|---|---|---|---|---|
| | Estimate | Est.Error | l-95% CI | u-95% CI |
| Intercept | 0.51 | 0.06 | 0.37 | 0.60 |
| Phenotype (AN-NSF) | 0.06 | 0.03 | -0.00 | 0.13 |
| **Table S8B: Spatial importance** | | | | |
| | Estimate | Est.Error | l-95% CI | u-95% CI |
| Intercept | 0.47 | 0.09 | 0.25 | 0.59 |
| **Phenotype (AE-SF)** | **0.13** | **0.05** | **0.03** | **0.24** |
| **Table S8C: Habitat importance** | | | | |
| | Estimate | Est.Error | l-95% CI | u-95% CI |
| Intercept | 0.46 | 0.05 | 0.34 | 0.55 |
| **Phenotype (AN-SF)** | -0.16 | 0.07 | -0.30 | -0.02 |
